# Phosphorylation-controlled cohesion of a nuclear condensate regulates mRNA retention

**DOI:** 10.1101/2023.08.21.554101

**Authors:** Alexa B. R. McIntyre, Adrian Beat Tschan, Katrina Meyer, Severin Walser, Arpan Kumar Rai, Keisuke Fujita, Lucas Pelkmans

**Author notes:** these authors contributed equally (Alexa B. R. McIntyre, Adrian Beat Tschan).

## Abstract

Nuclear speckles are membraneless organelles that associate with active transcription sites and participate in post-transcriptional mRNA processing. During the cell cycle, nuclear speckles dissolve following phosphorylation of their protein components. Here, we identify the PP1 family as the phosphatases that counteract kinase-mediated dissolution. PP1 overexpression increases speckle cohesion and leads to retention of polyadenylated RNA within speckles and the nucleus. Using APEX2 proximity labeling combined with RNA-sequencing, we characterized the relationship between the cohesion of nuclear speckles and the recruitment of specific RNAs. We find that many transcripts are preferentially enriched within nuclear speckles compared to the nucleoplasm, particularly chromatin- and nucleus-associated transcripts. While total polyadenylated RNA retention increased with nuclear speckle cohesion, the ratios of most mRNA species to each other were constant, indicating non-selective, or proportional, retention. We then explored whether nuclear speckle cohesion changes in response to environmental perturbations associated with changes in kinase or phosphatase activity. We found that cellular responses to heat shock, oxidative stress, and hypoxia include changes to the cohesion of nuclear speckles and mRNA retention. Our results demonstrate that tuning the material properties of nuclear speckles provides a mechanism for the acute control of mRNA localization.

## Introduction

Specific proteins and long non-coding RNAs form biomolecular condensates called nuclear speckles. In particular, the proteins SON and SRRM2 act as scaffolds for their formation, while the long non-coding RNA MALAT1 is highly enriched (Hutchinson et al., 2007; Ilık et al., 2020; Miyagawa et al., 2012). Splicing factors are proposed to cycle between speckles and active splicing sites in response to changes in their phosphorylation (Cao et al., 1997; Mermoud et al., 1994). Messenger RNAs (mRNAs) also localize to speckles and have been shown to rapidly exchange between nuclear speckles and the nucleoplasm (Politz et al., 2006). Recently, RNA sequencing after proximity-based labeling showed differential speckle association among transcripts (Barutcu et al., 2022). Although nuclear speckles lack the well-defined layered structure of nuclear condensates like nucleoli, the two scaffold proteins SON and SRRM2 tend to be more enriched within the centre of speckles, while MALAT1, mRNAs, and splicing factors tend towards the periphery (Fei et al., 2017). Within the nucleus, nuclear speckles associate with sites of active transcription and boost transcription of proximal genes (Hu et al., 2010; Kim et al., 2020; L. Zhang et al., 2021).

So far, few studies have explored how the morphology and material properties of nuclear speckles relate to their proposed functions. Nuclei contain around 20-40 nuclear speckles, with estimated areas of around 2 µm^2^ (Galganski et al., 2017; Q. Zhang et al., 2016). Inhibition of transcription and splicing have both been reported to increase the size of nuclear speckles (Kim et al., 2019; Misteli & Spector, 1996; O’Keefe et al., 1994; Rino et al., 2007). Transcriptional inhibition also increases their mobility within the nucleus and the exchange of splicing factors between speckles and the nucleoplasm (Kim et al., 2019; Rino et al., 2007). Recently, expression of a synthetic arginine-rich mixed-charge domain in cells was reported to increase the cohesion, or solidity, of nuclear speckles, based on a decrease in the exchange of proteins between speckles and the nucleoplasm, and to increase the enrichment of polyA FISH signal in the nucleus (Greig et al., 2020). The material properties of nuclear speckles could therefore be important in the control of mRNA localization, but it is unclear whether this mechanism of regulation is actively controlled under physiological conditions.

Nuclear speckles undergo a cycle of dissolution during early mitosis through the activity of kinases CLK1 and DYRK3. They then re-form between late metaphase and telophase (Colwill et al., 1996; Galganski et al., 2017; Rai et al., 2018). Most research has focused on the activity of kinases in the control of condensates, and we have little insight into the effects of dephosphorylation by phosphatases. In general, two families of serine-threonine phosphatases, PP1 and PP2A, control an estimated 90% of dephosphorylation events within mammalian cells (Bollen et al., 2010). The two families share some overlapping substrates; however, the catalytic phosphatases of each family also show sequence-based preferences (Hoermann et al., 2020) and gain additional specificity through association with regulatory subunits (Bertolotti, 2018). Both PP1 and PP2A are associated with functional splicing (Shi et al., 2006; Trinkle-Mulcahy et al., 1999), suggesting potential links to the regulation of nuclear speckles. In C. elegans, a PP2A phosphatase homolog was found to counteract the dissolution of P granules mediated by a DYRK-family kinase, MBK2 (J. T. Wang et al., 2014). However, which phosphatases counteract kinase activity to regulate nuclear speckles in mammalian cells is unknown.

Various perturbations can modulate the phosphorylation of nuclear speckle-associated proteins. For example, hypoxia results in increased CLK1 expression and phosphorylation of splicing factors (Jakubauskiene et al., 2015). On the other hand, phosphorylation of splicing factors decreases in response to heat shock, leading to differential splicing, export, and increased translation of the *CLK1* transcript, important for stress recovery (Haltenhof et al., 2020; Ninomiya et al., 2011).

Here, we hypothesized that differential phosphorylation of nuclear speckle-associated proteins affects the material properties of the structures, not only in mitotic cells during dissolution, but also in interphase cells to affect mRNA localization. Indeed, we found that overexpression of PP1 phosphatases or drug inhibition of DYRK3 or CLK1 increases nuclear speckle cohesion and mRNA enrichment in speckles. Sequencing of speckle-enriched RNAs suggested global, non-selective retention of mRNAs following dephosphorylation of nuclear speckles, with few genes disproportionately retained. Both heat shock and oxidative stress increased nuclear speckle cohesion and nuclear mRNA retention, consistent with lower phosphorylation, while hypoxia produced the opposite effects. Our results point to a new mechanism for rapid and dynamic control of mRNA localization through modulation of the material properties of an RNA-protein condensate.

## Results

### PP1 phosphatases counteract the effects of DYRK3/CLK1 and increase the cohesion of nuclear speckles

Nuclear speckles comprise protein and RNA components, including the scaffold proteins SON and SRRM2 and polyadenylated (polyA) RNA (**Figure 1A**). We first investigated which phosphatases can maintain nuclear speckle structure during kinase overexpression. For this, we overexpressed the kinase DYRK3, previously shown to dissolve nuclear speckles at high concentration during interphase (Rai et al., 2018), and co-overexpressed GFP-tagged phosphatases in HeLa cells. Overexpression of the PP1 catalytic subunit PPP1CC increased the concentration of DYRK3 necessary for speckle dissolution, an effect not seen with overexpression of a PP2A phosphatase (PPP2CA) (**Figure 1B**, **Supplementary Figure 1A**). Based on these results, we hypothesized that PPP1CC and DYRK3 share similar substrates within nuclear speckles.

**Figure 1:**
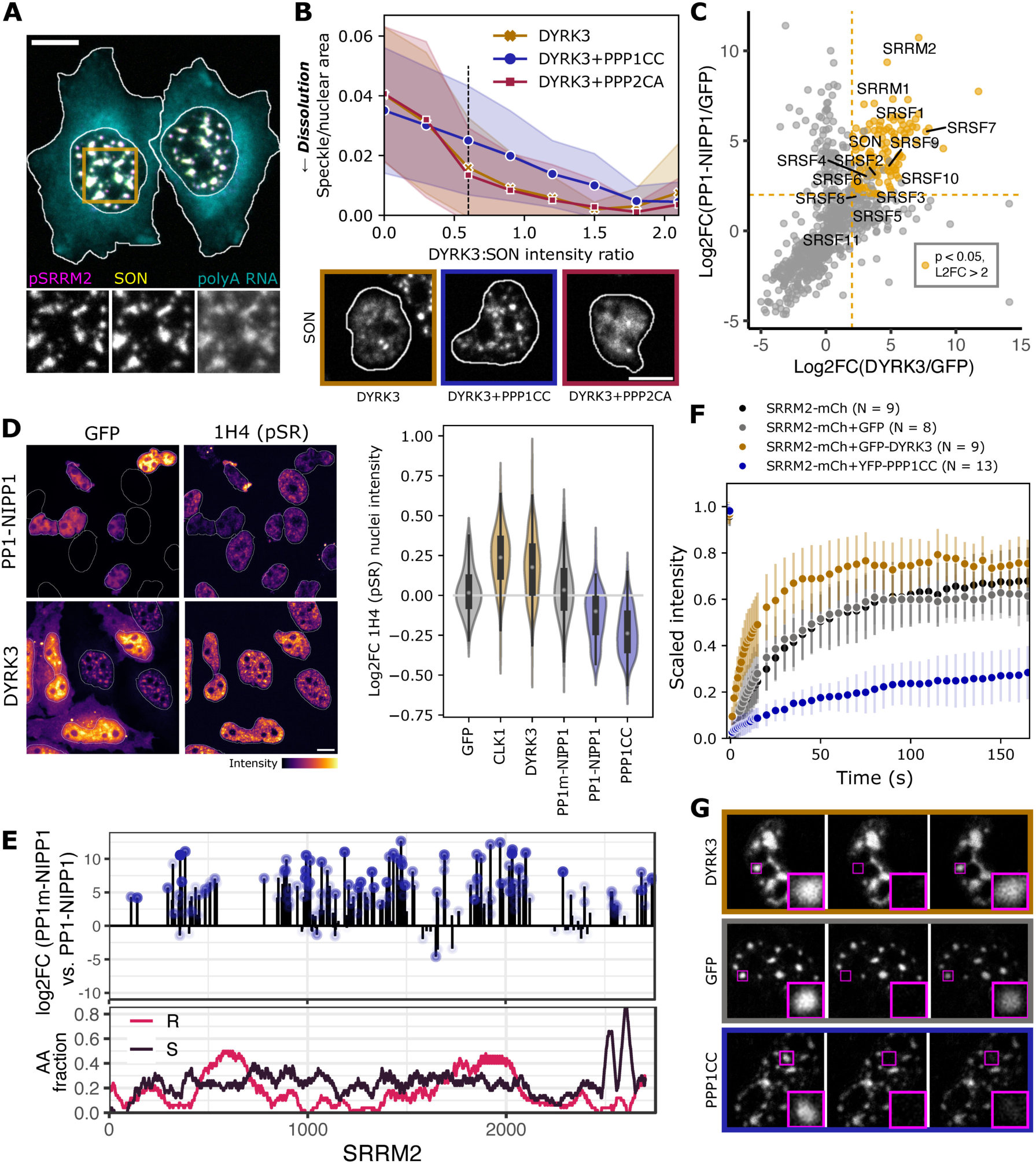
A phosphorylation cycle regulated by kinases DYRK3 and CLK1 and PP1 phosphatases controls the cohesion of nuclear speckles. **(A)** PolyA FISH and immunofluorescence imaging shows colocalization of mRNA and nuclear speckle proteins SON and SRRM2. **(B)** Above: overexpression of mCh-DYRK3 dissolves nuclear speckles, an effect counteracted by co-overexpression of GFP-PPP1CC but not GFP-PPP2CA. Shaded regions in line plot correspond to standard deviations. Below: staining for SON in example cells at the same level of mCh-DYRK3 expression (indicated by dotted line). Nuclear segmentations are shown for transfected cells. **(C)** Pulling down GFP-DYRK3 and GFP-PP1-NIPP1 enriches for similar protein interactors, including nuclear speckle scaffold proteins and splicing factors. Highlighted points indicate proteins pulled down with L2FC > 2 and p < 0.05 for both genes compared to GFP. **(D)** Overexpression of GFP-PP1-NIPP1 or YFP-PPP1CC reduces phospho-SR within the nucleus, while overexpression of GFP-DYRK3 or GFP-CLK1 increases phospho-SR. Example images are shown at left, coloured by relative pixel intensities. Quantification across nuclei is shown at right, with active kinases in yellow, active phosphatases in blue, and controls (GFP and mutant catalytic phosphatase) in grey (N > 70 cells per condition). Log2FC was calculated relative to the mean pSR intensity for untransfected cells in the same well. **(E)** The nuclear speckle scaffold protein SRRM2 shows differential phosphorylation depending on whether active (GFP-PP1-NIPP1) or mutant catalytic (GFP-PP1m-NIPP1) phosphatase is overexpressed. Shading corresponds to p-value (darker for lower p). Below: arginine (R) and serine (S) enrichment across the SRRM2 protein. **(F)** Fluorescence recovery after photobleaching of SRRM2-mCh shows increased recovery with GFP-DYRK3 overexpression and decreased recovery (a higher immobile fraction) with YFP-PPP1CC overexpression compared to controls. Error bars show standard deviations. **(G)** Examples of FRAP recovery under the conditions plotted above. Scale bars correspond to 10 µm.

Previous research established that the regulatory subunit NIPP1, or PPP1R8, retargets PP1 catalytic subunits from the nucleolus and elsewhere in the cell to nuclear speckles (Trinkle-Mulcahy et al., 2001) (**Supplementary Figure 1B**). We therefore used a GFP-tagged construct of the catalytic subunit PPP1CC fused to NIPP1 (PP1-NIPP1) to enrich for protein targets of PP1 within nuclear speckles. Following immunoprecipitation-mass spectrometry (IP-MS), we compared proteins enriched with IP of GFP-PP1-NIPP1 to those enriched with GFP-DYRK3. Many serine/arginine-rich (SR) splicing factors, as well as the nuclear speckle scaffold proteins SON and SRRM2, interacted with both phosphatase and kinase (**Figure 1C**). We used an antibody (1H4) that detects phosphoepitopes in serine/arginine-rich proteins to confirm that overexpression of either DYRK3 or CLK1 increases phosphorylation of SR regions (pSR) in splicing factors, while PP1-NIPP1 or PPP1CC overexpression decreases pSR in immunofluorescence imaging (**Figure 1D**, **Supplementary Figure 1C**). A catalytically impaired mutant version of PP1 joined to NIPP1 (GFP-PP1m-NIPP1 (Wu et al., 2018)) showed no effect on pSR.

To confirm whether PP1-NIPP1 overexpression also increases dephosphorylation of the scaffold proteins of nuclear speckles, we performed phosphoproteomics after IP of GFP-PP1-NIPP1 or GFP-PP1m-NIPP1. Many sites (148 with log2 fold change ≥ 2, p < 0.05) within SRRM2 showed differential phosphorylation depending on whether the fully active or mutant catalytic phosphatase was overexpressed (**Figure 1E**). We detected only five sites in the second scaffold protein of nuclear speckles, SON, that showed significant differences in phosphorylation depending on fully active or mutant phosphatase overexpression (Hornbeck et al., 2015) (**Supplementary Figure 1E**). Fifty PP1-regulated sites across proteins were also differentially phosphorylated by DYRK3 compared to DYRK3 inhibited by small compound inhibitor GSK626616, with 12 of those sites within SRRM2 **(Supplementary Figure 1D-E)**. However, we note that the dissolution of nuclear speckles in response to overexpressed DYRK3 compared to increased enrichment of DYRK3 in nuclear speckles upon inhibition (Rai et al., 2018) could affect which interactions we detect.

To then study the effects of phosphorylation on the material properties of nuclear speckles, we measured fluorescence recovery after photobleaching (FRAP) of mCherry-tagged SRRM2. Overexpression of DYRK3 decreased the cohesion of SRRM2 condensates and led to higher recovery from photobleaching, suggesting increased mobility of SRRM2. Meanwhile, overexpression of PPP1CC or its fellow PP1 catalytic subunits, PPP1CA and PPP1CB, increased the cohesion of SRRM2 and decreased recovery from photobleaching (**Figure 1F-G**, **Supplementary Figure 1F**). A PP2A family phosphatase did not show similar effects. Despite enrichment of PP1 phosphatases within the nucleolus in addition to nuclear speckles, PP1 overexpression did not affect the cohesion of nucleolin, showing that PP1-mediated regulation is not universal across condensates (**Supplementary Figure 1G**). Proteomics data, immunofluorescence staining, phosphoproteomics, and FRAP experiments thus establish that PPP1CC dephosphorylates nuclear speckle-associated proteins to counteract the effects of DYRK3 and increase the cohesion of the nuclear speckles.

### Interfering with the cohesion of nuclear speckles affects mRNA retention

To determine how changes in the cohesion of nuclear speckles mediated through phosphorylation could affect mRNA regulation, we stained for bulk mRNA using polyA FISH. Overexpression of PPP1CC increased the polyA signal within nuclear speckles and the ratio of nuclear-to-cytoplasmic polyA RNA, while a PP2A-family phosphatase did not (**Figure 2A-B**, **Supplementary Figure 2A-B**). By contrast, the dissolution of nuclear speckles through DYRK3 overexpression correlated with a decrease in nuclear polyA RNA (**Figure 2C**). Similar correlations were observed by dissolving speckles through overexpression of GFP-CLK1 or the cargo-binding domain of TNPO3, which interacts with SR-rich regions (Hochberg-Laufer et al., 2019) (**Supplementary Figure 2C**). Altogether, this implies that modulating the cohesion of nuclear speckles affects mRNA retention within the nucleus.

**Figure 2:**
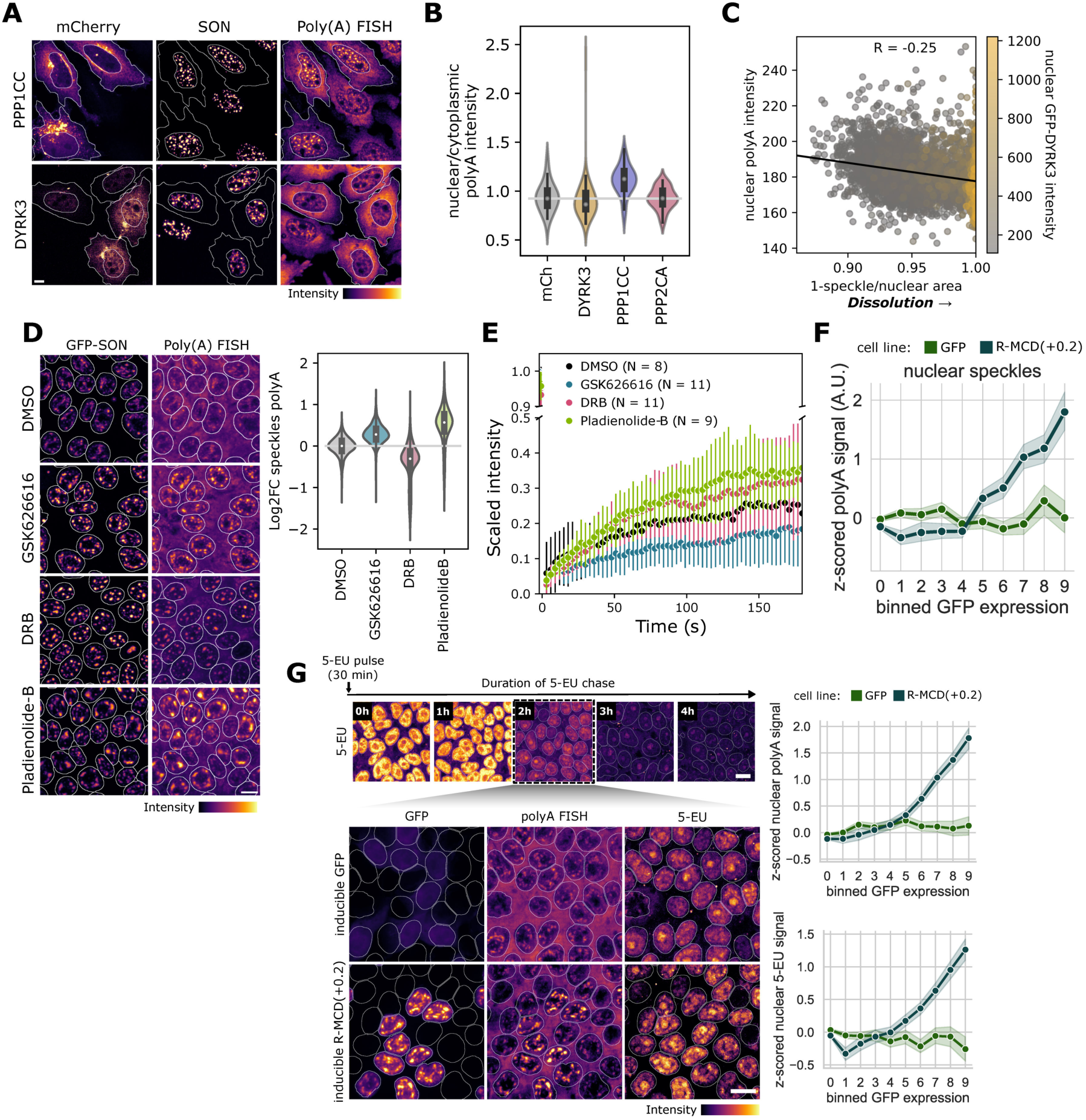
Perturbing nuclear speckle cohesion affects mRNA retention. **(A)** Example images showing differential polyA FISH intensity with PPP1CC and DYRK3 overexpression. mCherry images were scaled separately for PPP1CC and DYRK3, with outlines shown for transfected cells. SON and polyA FISH intensities were normalized based on untransfected cells in the same well. **(B)** Quantification for the ratio of nuclear to cytoplasmic polyA intensities for data in (A) and Supplementary Figure 2A. The top 5% of cells based on mCherry intensity were considered for each condition. **(C)** Nuclear polyA intensity negatively correlates with dissolution of nuclear speckles through GFP-DYRK3 overexpression. **(D)** Example images and quantification showing polyA intensity in nuclear speckles in hiPSCs under control conditions (DMSO), inhibition of DYRK3 (GSK626616), inhibition of transcription (DRB), and inhibition of splicing (Pladienolide-B). **(E)** Recovery from photobleaching in endogenously tagged 2xGFP-SON hiPSCs increased with transcriptional or splicing inhibition but decreased with DYRK3 inhibition. Error bars show standard deviations. **(F)** Overexpression of a positively charged arginine-rich mixed charge domain (R-MCD+02) leads to accumulation of polyadenylated RNA in nuclear speckles. **(G)** Nuclear PolyA FISH intensity and 5-EU nuclear intensity after 30 minutes of incubation with 5-EU and 2 hours of chase are elevated depending on the expression level of a positively charged arginine-rich mixed charge domain (R-MCD+0.2). Scale bars correspond to 10 µm.

We further measured an increase in polyA RNA signal within nuclear speckles by inhibiting DYRK3 using the small molecule inhibitor GSK626616 in human induced pluripotent stem cells (hiPSCs) expressing endogenously-tagged GFP-SON. Under DYRK3 inhibition, speckles became larger and rounder (**Figure 2D**). Previous reports described similar effects on the morphology of speckles under transcriptional or splicing inhibition (Kim et al., 2019; Misteli & Spector, 1996; O’Keefe et al., 1994; Rino et al., 2007). We therefore compared DYRK3 inhibition to the inhibition of transcription using DRB and the inhibition of splicing using Pladienolide-B to determine whether all exerted similar effects on nuclear speckle cohesion and RNA regulation. PolyA RNA retention in nuclear speckles increased with DYRK3 inhibition and splicing inhibition, but decreased under transcriptional inhibition, likely due to ongoing mRNA export. The nucleus as a whole similarly showed an increase in polyA retention upon DYRK3 or splicing inhibition, but we measured no change within the nucleoplasm (**Supplementary Figure 2D**). Meanwhile, the cohesion of nuclear speckles increased under DYRK3 inhibition, as measured using FRAP, but decreased in response to transcriptional or splicing inhibition (**Figure 2E**). Although GSK626616 and Pladienolide-B share effects on the morphology of nuclear speckles as well as on polyA RNA retention, the material properties of the speckles differ under the two inhibitors. Importantly, these differences suggest that increased nuclear mRNA retention (as with splicing inhibition) does not increase nuclear speckle cohesion.

We next generated an inducible hiPSC line for phosphorylation-independent modulation of nuclear speckle cohesion. Expressing a positively charged arginine-rich mixed charge domain (R-MCD+0.2) in hiPSCs increased nuclear and nuclear speckle mRNA intensity, as previously reported (Greig et al., 2020) (**Figure 2F-G**, **Supplementary Figure 2E**). We also observed an increase of polyA FISH signal in the nucleoplasm, but no change in the cytoplasm (**Supplementary Figure 2E**). To further verify that the observed increase in nuclear intensity represents transcript retention, we then used 5-EU to label nascent transcripts. A 30-minute pulse of 5-EU incorporation, followed by a 2-hour chase showed that newly generated mRNA remains within the nucleus for an extended time when cohesion increases through R-MCD+0.2 expression (**Figure 2G**). Notably, R-MCD+0.2 overexpression led to a decrease in 5-EU incorporation (**Supplementary Figure 2F**), as previously reported, consistent with feedback between nuclear retention of mRNA and transcription rate (Berry et al., 2022). Thus, increasing the cohesion of nuclear speckles either through dephosphorylation or expression of a mixed charge domain increases bulk mRNA retention in the nucleus.

### Characterization of the nuclear speckle transcriptome by APEX2-seq

To explore if bulk enrichment of polyA RNA upon increased cohesion reflected changes in nuclear speckle RNA composition, we characterized the nuclear speckle transcriptome. Until recently, the localization of specific mRNAs to nuclear speckles had been characterized for only a few genes (Bahar Halpern et al., 2015; Smith et al., 1999; K. Wang et al., 2018). Proximity-based labelling followed by RNA sequencing has now enabled transcriptomic profiling of distinct subcellular compartments (Fazal et al., 2019; Padrón et al., 2019), including nuclear speckles (Barutcu et al., 2022).

We used a proximity-based labelling approach to characterize the nuclear speckle transcriptome of hiPSCs. To avoid the effects of plasmid overexpression, we used CRISPR/Cas9 to tag the endogenous nuclear speckle scaffold protein SON with the engineered ascorbate peroxidase APEX2. As a control, a second cell line was generated by inserting APEX2 fused to a nuclear localization signal (NLS) into the genome at a safe-harbour locus to assess the overall nuclear transcriptome (**Figure 3A**). After inducing the APEX2 enzymatic activity by addition of biotin-phenol and hydrogen peroxide, we detected biotinylated RNA using a dot-blot assay (**Supplementary Figure 3A**). Staining for biotin using fluorescent streptavidin showed distinct localization of the streptavidin signal to nuclear speckles in the APEX2-SON cell line, and to the nucleus in the APEX2-NLS cell line (**Figure 3B**). We collected RNA from both cell lines following 2-hour treatment with DYRK3 inhibitor (GSK626616) or DMSO (control) to evaluate changes in the nuclear speckle transcriptome in response to increased nuclear speckle cohesion. We also included treatments with DRB and Pladienolide-B to compare the effect of increased nuclear speckle cohesion to transcriptional inhibition and splicing inhibition (**Figure 3C**).

**Figure 3:**
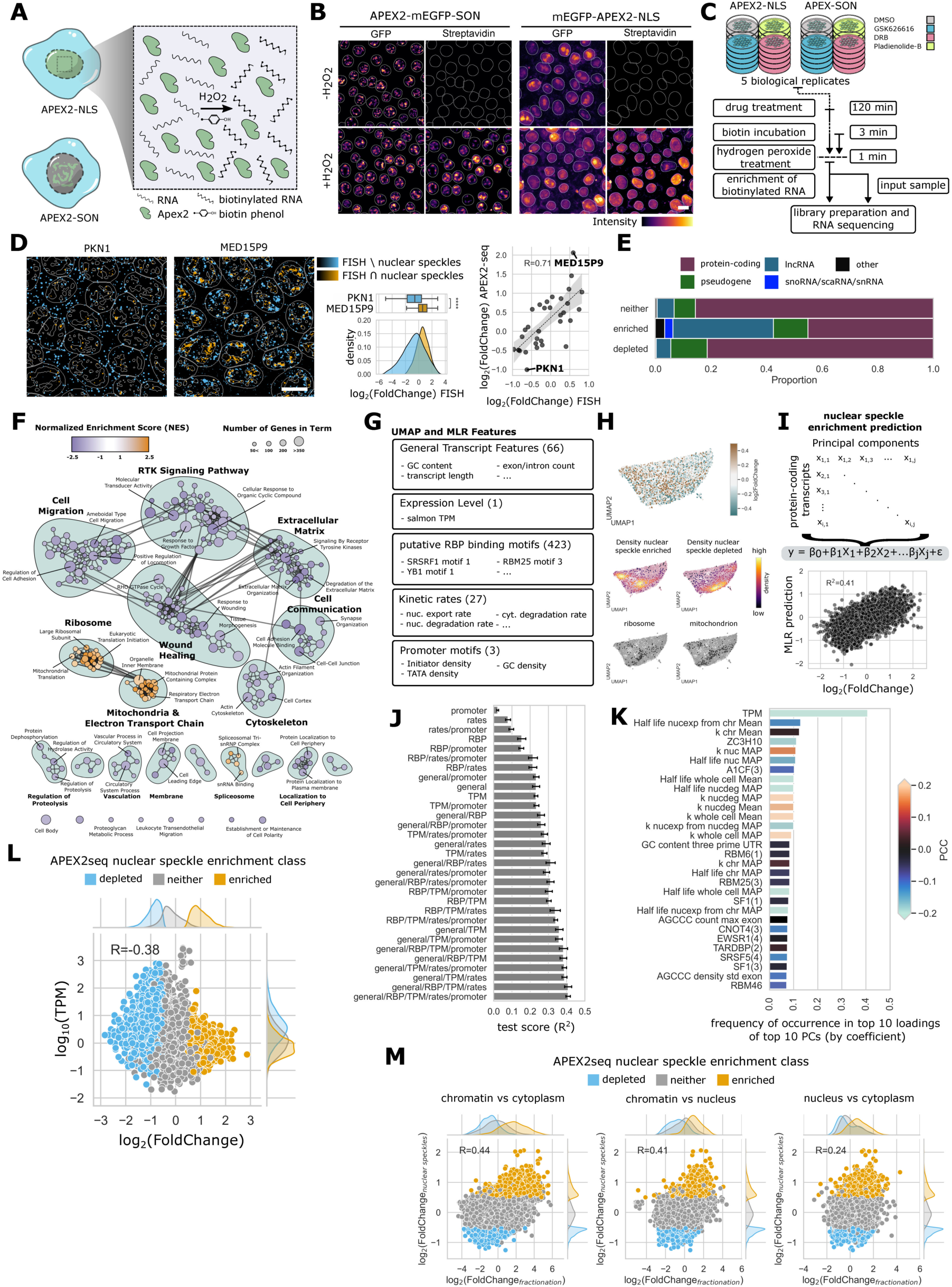
Characterization of the nuclear speckle transcriptome by APEX2-seq. **(A)** Schematic illustration of APEX2-NLS and APEX2-SON human induced pluripotent stem cells and APEX2 enzymatic reaction leading to biotinylation of RNA. **(B)** Activation of APEX2 enzymatic activity by addition of hydrogen peroxide leads to biotinylation of biomolecules near nuclear speckles in APEX2-SON hiPSCs (left) and in the nucleoplasm in APEX2-NLS hiPSCs (right). Scale bar corresponds to 10 µm. **(C)** APEX2-seq experimental outline. APEX2-NLS and APEX2-SON hiPSCs were treated with GSK626616, DRB or Pladienolide-B for two hours prior to addition of biotin-phenol and activation of APEX2-enzymatic activation by hydrogen peroxide. 5 biological replicates were collected per condition. **(D)** Example pseudocolor images of smFISH signal for two gene targets that were found to be nuclear speckle depleted (PKN1) or enriched (MED15P9) in DMSO control conditions. An orange colormap was used to map FISH signal overlapping with nuclear speckles (FISH ∩ nuclear speckles), while a cyan colormap was used to map FISH signal that does not overlap with nuclear speckles (FISH \ nuclear speckles). Outlines of nuclei and nuclear speckles are indicated by white overlays. Density plots show single cell log2FoldChange of nuclear speckle vs. nucleoplasm FISH signal. Boxplots of the same data are shown with an indicator for the median. Asterisks indicate significance based on one-sided Wilcoxon signed-rank tests (****p <= 0.0001). APEX2-seq and smFISH show positive correlation (Pearson correlation coefficient = 0.71). Scale bar corresponds to 10 µm. **(E)** Composition of nuclear speckle transcriptome. Nuclear speckle enriched subset of genes displays a larger proportion of long non-coding RNA and snoRNA/scaRNA/snRNA than nuclear speckle depleted and neither subsets. **(F)** Network representation of Gene Set Enrichment Analysis result. Terms associated with ribosome, mitochondria and electron transport chain and the spliceosome are enriched in nuclear speckles. **(G)** Feature categories that were used to build UMAP and train the multilinear regression model. Number of features in each category are indicated in brackets. Example features are displayed for each category. **(H)** UMAP representation of the multidimensional single transcript feature space. Nuclear speckle depleted/enriched transcripts occupy distinct regions of this feature space. Gene sets that were found to be enriched in nuclear speckles by GSEA show similar distributions. **(I)** Features from (G) were used to train a multilinear regression model to predict nuclear speckle enrichment of protein coding genes on a single-transcript level. Scatterplot shows random samples of 600 predictions/measurement pairs on test sets aggregated over 10-fold cross validation. **(J)** Training the multilinear regression model on all possible combinations of feature categories. Error bars indicate the standard deviation of ten models. **(K)** Frequency of feature occurrence in top ten loadings of the top ten PCs (by coefficient) of 100 models. A value of 1 indicates that the feature appears in the top loadings of each of the top 10 PCs. Color indicates Pearson correlation with transcript nuclear speckle enrichment. **(L)** TPM (transcripts per million) and nuclear speckle enrichment of protein coding transcripts show a negative correlation (Pearson correlation coefficient = −0.38) **(M)** Comparison of nuclear speckle enrichment based APEX2-seq and enrichment in different subcellular compartments based on fractionation sequencing. Nuclear speckle enrichment shows positive correlation with enrichment in the chromatin fraction over the nucleus and the cytoplasm, and with enrichment in the nucleus fraction over the cytoplasm fraction.

We started by analysing baseline transcript enrichment in nuclear speckles under control conditions. Differential gene expression analysis revealed 878 genes that showed significant enrichment and 713 genes that were depleted from nuclear speckles (|log2 fold change| > 0.5 and padj < 0.05) (**Supplementary Figure 3B**). We then designed single-molecule FISH (smFISH) probes against a subset of hits for validation (**Supplementary Figure 3C**). Enrichment measured by smFISH (quantified as the ratio of mean FISH signal in nuclear speckles over the remaining nucleoplasm) correlated well with enrichment detected by APEX2 sequencing (Pearson correlation coefficient = 0.71) (**Figure 3D**). Based on sequencing, genes enriched in nuclear speckles included a higher proportion of long non-coding RNAs and a category of RNAs including small nuclear RNAs (snRNAs), small Cajal body-specific RNAs (scaRNAs) and small nucleolar RNAs (snoRNAs) (**Figure 3E**). Gene set enrichment analysis showed that genes enriched in nuclear speckles were associated with ribosomal proteins and oxidative phosphorylation, whereas genes depleted from nuclear speckles were associated with a diverse set of terms, among them cytoskeletal and cytoplasmic localization terms (**Figure 3F**).

We next compared our APEX2-seq dataset to the previously published proximity labeling dataset from Barutcu et al., 2022. In this study, nuclear speckle-enriched genes were identified by expressing APEX2 fused to three splicing factors in HEK293T cells. We found overlaps of 354, 436, and 268 for genes enriched with SRSF1, SRSF7, and RNSP1, respectively (**Supplementary Figure 3D**). Our pairwise comparisons of gene sets enriched or depleted in the SRSF1, SRSF7 and RNSP1 datasets showed an average Jaccard index of 0.226 between the different markers, whereas we found an average Jaccard index of 0.077 for the three markers compared to our dataset. This indicates that the overlap between RNAs in proximity with different markers for nuclear speckles is relatively low, and that there might be substantial differences in nuclear speckle RNA composition between different cell types.

As suggested by previous literature (Barutcu et al., 2022; Dias et al., 2010; Wegener & Müller-McNicoll, 2018), intron retention was higher in nuclear speckle-associated mRNA compared to the nucleus, with at least twice as many retained introns detected in speckles compared to the nucleus depending on the splicing analysis tool used (**Supplementary Figure 3E**). Consistent with the results of Barutcu et al., 2022, introns retained in speckles also showed higher GC content (mean ~49% vs. ~45% for randomly selected introns). However, differences in intron length between introns retained in speckles and the nucleus were inconsistent across splicing analysis tools (**Supplementary Figure 3F**).

Next, we aimed to predict nuclear speckle enrichment of single transcripts. We performed differential transcript enrichment analysis and bioinformatically integrated transcript features from various sources. These features included general sequence features (e.g., GC content, number of exons), transcript expression level (transcripts per million, TPM), putative RNA binding protein (RBP) binding motifs, kinetic rates, and promoter motifs (**Figure 3G, Supplementary Table 3**). Among these features, we find exon length positively correlated with nuclear speckle enrichment, while transcript expression level (TPM) anti-correlated with nuclear speckle enrichment (**Supplementary Figure 3G**). Embedding these features into a uniform manifold approximation and projection (UMAP) (McInnes, Healy, & Melville, 2018; McInnes, Healy, Saul, et al., 2018) reveals that transcripts enriched in or depleted from speckles occupy distinct regions within this feature space. Genes belonging to the enriched GO categories associated with the ribosome and mitochondria display similar distribution patterns, suggesting that transcript similarities may influence their association with nuclear speckles (**Figure 3H**).

To assess the predictive power of these features, we trained a multilinear regression model to predict nuclear speckle enrichment on a single transcript level. Using this approach, we achieve an average coefficient of determination (R^2^) of 0.41 (± 0.008, 95% CI) (**Figure 3I/J**). We trained the model on all possible combinations of feature categories. A model trained on minimal sets of general sequence features, transcript expression level and either RBP motif (R^2^ = 0.38 ± 0.0123) or rate features (R^2^ = 0.39 ± 0.0092) performed similarly to a model incorporating all the feature sets, suggesting that these are the most determining factors in speckle localization (**Figure 3J**). A model based on sequence features alone (general, RBP, and promoter motifs) reached an average R^2^ of 0.26 (± 0.013), while a model based solely on transcript expression level reached an average R^2^ of 0.23 (± 0.0066).

We then measured the importance of individual features by training multiple iterations of the model and calculating the frequency with which a feature appeared in the top loadings of the top principal components. By a large margin, transcript expression level (TPM) appears as the most important feature in this analysis (**Figure 3K**, **Supplementary Figure 3G**). Interestingly, although expression level showed an inverse correlation with nuclear speckle enrichment (**Figure 3K-L**), extending this analysis to all subtypes of RNA revealed a subset of highly expressed nuclear speckle-enriched transcripts composed mainly of snoRNAs, scaRNAs, and snRNAs (**Supplementary Figure 3H**).

To assess whether enrichment of transcripts in nuclear speckles relative to the nucleus correlates with enrichment of transcripts in the nucleus relative to the cytoplasm, we biochemically fractionated hiPSCs into chromatin-associated nucleus, nucleus, and cytoplasm fractions and performed RNA sequencing. We found that nuclear speckle-associated transcripts tend to be enriched in the chromatin fraction compared to the nucleus and cytoplasm fractions, and to a lesser extent, in the nucleus compared to the cytoplasm (Pearson correlation coefficient of 0.41, 0.44 and 0.24, respectively) (**Figure 3M**). We likewise found a positive correlation between nuclear speckle enrichment and nuclear enrichment (quantified as ratio of nuclear over cytoplasmic FISH signal) in our smFISH dataset (Pearson correlation coefficient = 0.65, **Supplementary Figure 3I**).

Collectively, these results offer a comprehensive overview of transcript localization relative to nuclear speckles in hiPSCs. Statistical modeling showed that transcript expression level is a strong determinant of nuclear speckle enrichment. Contrary to sequence features, transcript expression level is a variant feature, suggesting that nuclear speckle enrichment of transcripts could differ between cell types and tissues. The general enrichment of speckle-associated transcripts within the nuclear fraction is consistent with a role for speckles in storing nuclear-detained mRNAs. Interestingly, this may be more important for transcripts that are less highly expressed, suggesting tighter buffering of their translational availability.

### An increase in nuclear speckle cohesion leads to proportional enrichment of mRNA

We next investigated whether the increased recruitment of bulk polyadenylated RNA we observed with increased speckle cohesion (see **Figure 2A-D**) reflects the differential retention of specific transcripts. We therefore compared APEX2 sequencing results of DYRK3-inhibited cells to our DMSO-treated controls. Surprisingly, few genes (N = 24) significantly changed in speckle enrichment under DYRK3 inhibition relative to other genes, whereas 759 and 1031 genes changed under transcriptional and splicing inhibition, respectively (log2 fold change| > 0.5, padj < 0.1) (**Figure 4A**). To reconcile this finding with the bulk enrichment of polyadenylated RNA, we hypothesized that enrichment of RNA is non-specific, and that most genes are retained proportionally in nuclear speckles upon increased cohesion (**Figure 4B**). Here, we did not include spike-in controls, as the biotinylation reaction and enrichment of biotinylated RNA could introduce technical variability between experiments prior to spike-in addition. We were therefore unable to quantify absolute changes in transcript abundances between samples from our APEX2 sequencing data. We note that generation and addition of a biotinylated spike-in before enrichment could help quantification in future experiments, although would still not control for variability from an initial biotinylation reaction.

**Figure 4:**
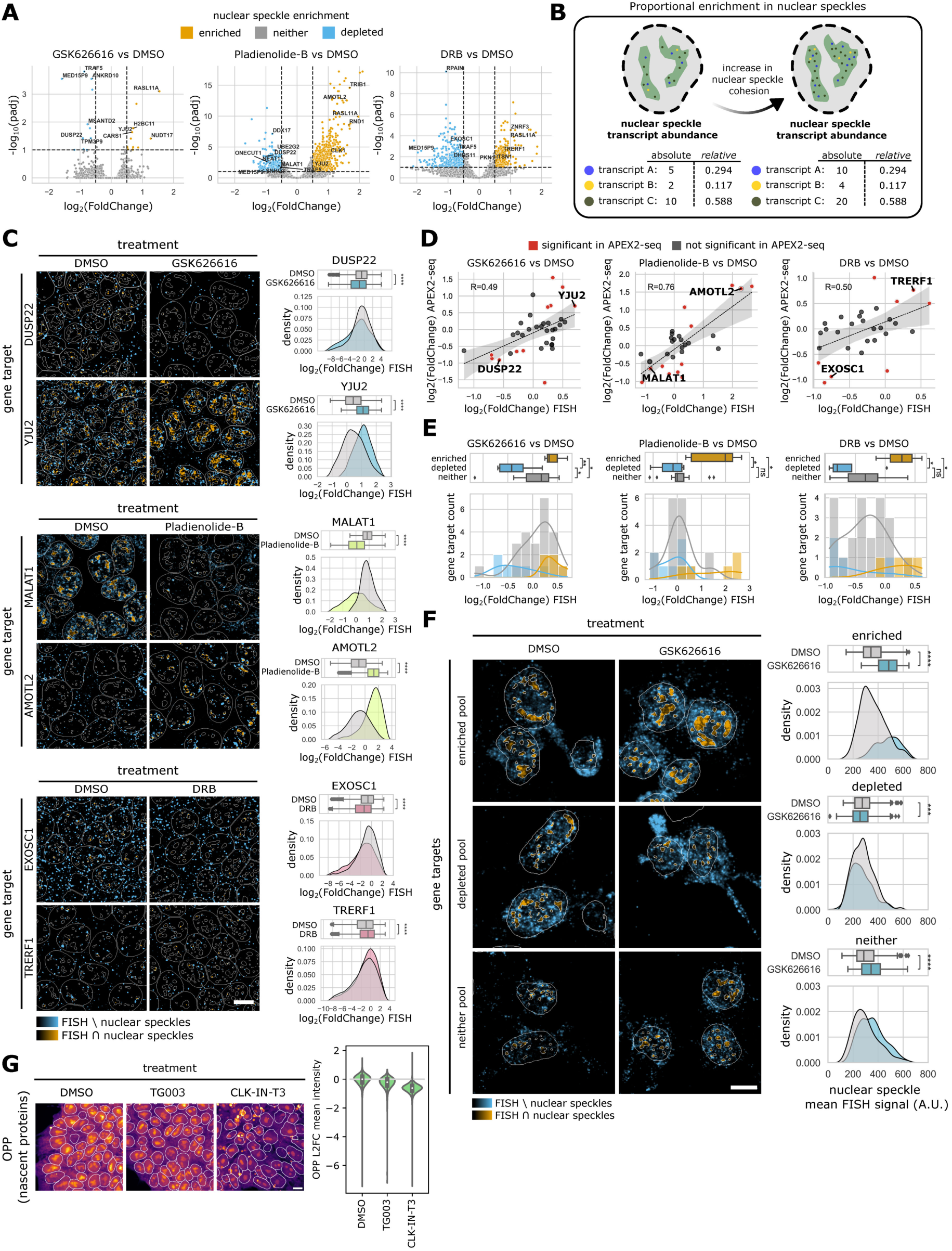
An increase in nuclear speckle cohesion leads to proportional enrichment of mRNA. **(A)** Volcano plots showing differentially expressed genes in nuclear speckles vs. nucleoplasm in GSK626616 vs. DMSO condition (left), Pladienolide vs. DMSO condition (centre) and DRB vs. DMSO condition (right). Annotated points highlight genes for which smFISH probes were designed. **(B)** Schematic illustration of proportional enrichment of RNA in nuclear speckles upon increase in speckle cohesion. **(C)** Example pseudocolor images of FISH signal for two gene targets that were found to be nuclear speckle depleted or enriched in each of the three drug conditions. DUSP22 (depleted) and YJU2 (enriched) in the GSK626616 vs. DMSO condition (left), MALAT1 (depleted) and AMOTL2 (enriched) in the Pladienolide-B bs. DMSO condition (centre), and EXOSC1 (depleted) and TRERF1 (enriched) in the DRB vs. DMSO condition (right). Outlines of nuclei and nuclear speckles are indicated by white overlays. Density plots show single cell log2FoldChange of nuclear speckle vs. nucleoplasm FISH signal. Boxplots of the same data are shown with an indicator for the median. Asterisks indicate significance based on one-sided Wilcoxon signed-rank tests (****p <= 0.0001). Scale bar corresponds to 10 µm. **(D)** Correlation of nuclear speckle enrichment in response to drug treatment in APEX2-seq vs. smFISH. Red dots highlight genes that were found to significantly change (|log2FoldChange| > 0.5, padj < 0.1) in APEX2-seq. Pearson correlation coefficient (calculated over all data points) is displayed. **(E)** Distributions of FISH Δdrug vs. DMSO nuclear speckle log2FoldChange. Colors indicate genes identified to be nuclear speckle enriched, depleted or neither upon GSK626616, Pladienolide-B or DRB treatment based on APEX2-seq. Boxplots of the same data are shown with an indicator for the median. Asterisks indicate significance based on Welch’s t-test (0.01 < *p <= 0.05, 0.001 < **p <= 0.01). Genes belonging to the “neither” class display a bias towards being more enriched in nuclear speckles upon GSK626616 treatment in smFISH. **(F)** Example pseudocolor images of FISH signal of pooled probe sets (n**≈**50 targets per pool) representing gene targets that were found to be nuclear speckle depleted or enriched or neither upon GSK treatment. Density plots show single cell mean nuclear speckle FISH signal. Boxplots of the same data are shown with an indicator for the median. Asterisks indicate significance based on one-sided Wilcoxon signed-rank tests (****p <= 0.0001, 0.0001 < ***p <= 0.001). Scale bar corresponds to 10 µm. **(G)** Example images and quantification across cells showing changes in translational rates in response to two inhibitors of the nuclear kinase CLK1, based on incorporation of the puromycin analog O-Propargyl Puromycin (OPP) into nascent peptides.

To test proportional enrichment, we used smFISH to quantify absolute RNA levels across compartments. As for baseline speckle enrichment, we found a positive correlation (Pearson correlation of 0.49-0.76) between APEX2 sequencing and smFISH (**Figure 4C-D**, **Supplementary Figure 4A-C**). Under DYRK3 inhibition, we observed a median log2FoldChange in nuclear speckle enrichment of 0.184 for gene targets that did not show disproportionate changes based on APEX2 sequencing. Inhibition of transcription or splicing did not result in enrichment of these gene targets in nuclear speckles (**Figure 4E**). This indicates a small general shift into nuclear speckles upon increased cohesion but not splicing inhibition. To further explore this, we designed three smFISH pools, each targeting around 50 random genes identified as enriched, depleted or left unchanged (neither) upon DYRK3 inhibition based on APEX2 sequencing results. Because few genes changed significantly when using more conservative thresholds, we defined a relaxed threshold of significance for enriched and depleted pools (|log2 fold change| > 0.3, padj < 0.6) and a very stringent threshold for the neither pool (|log2 fold change| < 0.05, padj > 0.99) (**Supplementary Figure 4D**). As anticipated, the enriched and depleted smFISH pools showed an increase (log2 fold change=0.5) and decrease (log2 fold change=-0.1), respectively, in localization of signal to nuclear speckles upon DYRK3 inhibition. However, the smFISH pool representing gene targets that showed no disproportionate changes in enrichment in APEX2 sequencing showed increased smFISH signal in nuclear speckles (log2 fold change=0.27) (**Figure 4F**). Notably, treatment with CLK-IN-T3, a CLK inhibitor, led to a similar enrichment of polyadenylated RNA in nuclear speckles as treatment with GSK626616, and we also observed non-specific enrichment when performing FISH with mixed smFISH pools (**Supplementary Figure 4F-G**). This further supports the notion of a general shift of the nuclear RNA population into nuclear speckles upon increasing speckle cohesion.

Previous work has also described an association between higher retention of transcripts within speckles and intron retention within transcripts (Girard et al., 2012; Martins et al., 2011). Changes in the association between specific transcripts and nuclear speckles could therefore result from decreased splicing efficiency under drug perturbations. Based on splicing analysis of the RNA-seq data, we found that a substantial subset (175/711) of genes more enriched in speckles in response to splicing inhibition also showed measurable increases in intron retention (**Supplementary Figure 4E**). With GSK626616, however, an order of magnitude more genes (425) showed changes in intron retention than disproportionate changes in nuclear speckle association (24, |log2 fold change| > 0.5, p < 0.1). The relatively few and bi-directional splicing changes that we find in response to DYRK3 inhibition are thus a poor explanation for broad retention of transcripts in speckles. Thus, while our results under baseline conditions and splicing inhibition are consistent with a role for intron retention in the recruitment of specific transcripts to nuclear speckles, splicing analysis suggests the global shift in transcript retention under DYRK3 inhibition is not driven by perturbed splicing.

Increased nuclear retention of transcripts implies decreased translational availability. Single-molecule FISH data shows that under DYRK3 kinase inhibition, the cytoplasmic concentration of most transcripts decreases and nuclear-to-cytoplasmic ratio increases (**Supplementary Figure 4H**). We further verified the effects of nuclear phosphorylation perturbations on translation. Inhibition of the nuclear-localized kinase, CLK1, led to a decrease in overall translation rates compared to controls (**Figure 4G**). Inhibition of DYRK3 did not decrease overall translation in this assay, although we note that cytoplasmic DYRK3 is known to regulate the mTOR pathway (Wippich et al., 2013) (**Supplementary Figure 4I**).

Taken together, our data supports a model in which RNA localizes to nuclear speckles relative to nuclear speckle cohesion. Our APEX2 sequencing results indicate that this relationship is rarely gene-specific but instead affects a large fraction of the nuclear RNA population equally, leading to proportional changes in enrichment of different RNAs as speckles become more solid-like. Measuring translation rates suggests that increased retention of mRNA in the nucleus correlates with decreased translation.

### Heat shock, oxidative stress, and hypoxia modulate nuclear speckle phosphorylation and mRNA retention

To explore a physiological role for the material properties of nuclear speckles, we asked whether environmental perturbations could affect mRNA retention in speckles through changes in phosphorylation. Previous research established that CLK1 expression and SR phosphorylation change in response to heat shock and hypoxia (Jakubauskiene et al., 2015; Ninomiya et al., 2011; Shin et al., 2004). We first exposed hiPSCs to these conditions to confirm measurable differences in SR phosphorylation in immunofluorescence. Heat shock (43°C for 1 hour) decreased SR phosphorylation, while hypoxia (0.2% oxygen for 24 hours) had the opposite effect. We further considered oxidative stress (0.5 mM sodium arsenite for 1 hour), which decreased pSR, similar to heat shock (**Figure 5A**, **Supplementary Figure 5A**). Using FRAP, we found that conditions in which SR phosphorylation decreased also showed higher cohesion of nuclear speckles, while hypoxia led to lower cohesion (**Figure 5B**, **Supplementary Figure 5B**).

**Figure 5:**
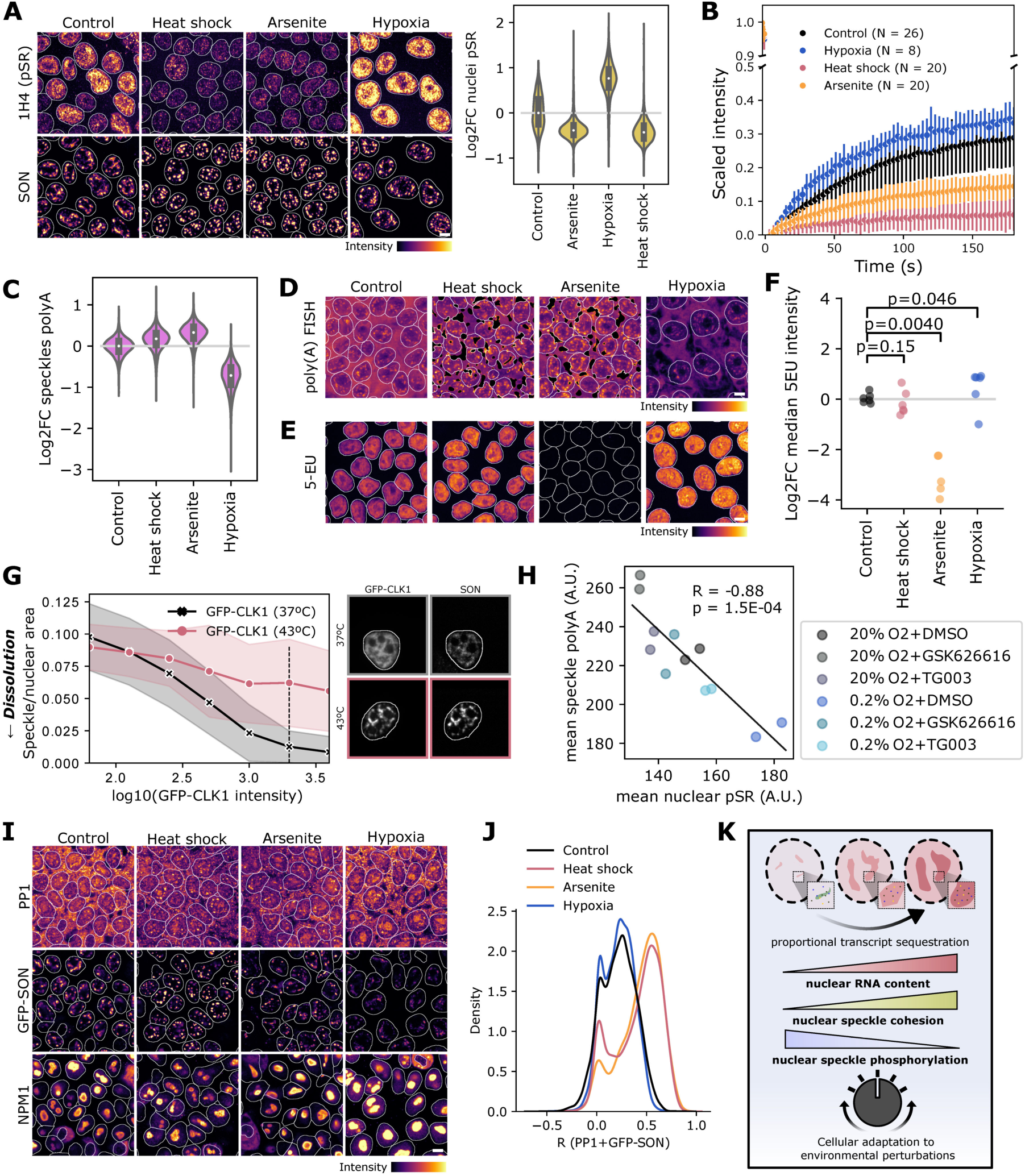
Nuclear speckle cohesion and nuclear speckle mRNA retention are adjusted in response to environmental perturbations. **(A)** Phosphorylation of SR sites decreases in the nucleus under heat shock (43°C, 1 hour) or arsenite treatment (0.5 mM, 1 hour) and increases with hypoxia (0.2% O2, 24 hours). **(B)** Fluorescence recovery of 2xGFP-SON-tagged nuclear speckles decreases under heat shock and arsenite treatment and increases under hypoxia compared to control. Error bars show standard deviations. **(C)** PolyA RNA retention in nuclear speckles increases with heat shock and arsenite treatment but decreases with hypoxia. **(D)** Example images of polyA FISH data quantified in (C). Stress granules were segmented out using a pixel classifier and are here shown in black. **(E)** Example images of 5-EU staining showing transcription levels are not altered by heat shock, decrease with arsenite stress, and increase under hypoxia. **(F)** Quantification of median nuclear 5-EU signal for two replicate wells in three experiments under different perturbations (p-values calculated by Mann–Whitney U test). **(G)** Dissolution of nuclear speckles by GFP-CLK1 overexpression is reversed by heat shock. Shaded regions in line plot correspond to standard deviations. **(H)** Mean nuclear pSR and speckle polyA FISH intensities across cells in each imaged well reveal an inverse correlation between phosphorylation (modulated by DYRK3 and CLK1 inhibitors) and speckle polyA RNA under hypoxia and control conditions. A.U. = arbitrary units. **(I)** Example images of PP1, GFP-SON, and NPM1 for data in (H) and Supplementary Figure 5F. **(J)** PP1 and GFP-SON pixel correlations across nuclei under different conditions. Scale bars correspond to 10 µM. **(K)** Schematic representation of the relationship between nuclear speckle phosphorylation, cohesion, and mRNA retention.

These changes in cohesion extended to the retention of polyadenylated RNA. Hypoxia decreased polyA FISH signal in both nuclear speckles and the cell as a whole. Heat shock and oxidative stress increased the polyA FISH signal within nuclear speckles, and to a lesser extent, the nucleoplasm. We also observed mRNA accumulation within stress granules in the cytoplasm under heat shock and oxidative stress (**Figure 5C-D**, **Supplementary Figure 5C-F**). Notably, changes in nuclear speckle retention did not reflect the effects of changed transcription rates based on 5-EU staining (**Figure 5E-F**, **Supplementary Figure 5G**). Oxidative stress inhibited RNA production and heat shock did not significantly affect transcription in iPSCs, as previously reported in HeLa cells (Zhang and Kleiner, 2019). By contrast, hypoxia increased transcription. Taken together, our data suggests that differential retention of mRNA under these conditions depends on nuclear speckle properties.

We then investigated whether we could rescue the observed changes in mRNA retention by modulating nuclear speckle cohesion. Similar to the prevention of DYRK3-mediated speckle dissolution by PPP1CC (**see Figure 1b**), 1 hour of heat shock reversed the dissolution of speckles normally observed with high levels of GFP-CLK1 overexpression (**Figure 5G**). However, in cells where dissolution did occur, heat shock failed to increase the nuclear retention of polyA RNA (**Supplementary Figure 5H**). Treating hiPSCs kept under hypoxic conditions for 24 hours with either CLK1 or DYRK3 inhibitors showed recovery of nuclear speckle polyA RNA levels after 2 hours of treatment (**Supplementary Figure 5I-J**). Mean nuclear pSR and mean speckle polyA RNA showed a strong negative correlation (R = −0.88, p = 1.5×10^−4^) across drug treatments under hypoxia and 20% O_2_ (**Figure 5H**). Similar to kinase inhibition, R-MCD+0.2 overexpression also restored nuclear speckle polyA RNA levels during hypoxia, showing that increasing cohesion is sufficient for the rescue of RNA retention (**Supplementary Figure 5K**).

Previous reports find changes in CLK1 expression under environmental perturbations (Jakubauskiene et al., 2015; Ninomiya et al., 2011; Shin et al., 2004), leaving unexamined a role for phosphatase availability. Although the involvement of regulatory subunits remains to be further elucidated, we observed enrichment of PP1 in nuclear speckles relative to the rest of the nucleus during oxidative stress and heat shock, but higher correlation with a nucleolar marker under hypoxia (**Figure 5I-J**, **Supplementary Figure 5L**). Overall, our results suggest that differences in mRNA localization in response to environmental perturbations can be regulated through changes in the cohesion of nuclear speckles, modulated through differential kinase and phosphatase activity at nuclear speckles.

Two of the conditions we considered (oxidative stress and heat shock) also induced stress granules. Stress granules are membraneless organelles, like nuclear speckles, and show accumulation of mRNAs during stress (Kedersha & Anderson, 2002). To determine whether similar transcripts are recruited to stress granules as retained in nuclear speckles, we next compared the nuclear speckle transcriptome to previously published data on the stress granule transcriptome (Khong et al., 2017). We found that these two gene sets were largely mutually exclusive: 120 genes were depleted from nuclear speckles but enriched in stress granules, and 74 genes were enriched in nuclear speckles but depleted from stress granules (**Supplementary Figure 5M**). By contrast, only three genes were enriched in both compartments and only 12 depleted from both. This mutual exclusivity could be related to the nuclear or cytoplasmic localization of transcripts and suggests a model wherein more nuclear transcripts tend to be retained within nuclear speckles, while more cytoplasmic transcripts are retained within stress granules in response to stress.

## Discussion

Here, we examine a possible role for nuclear speckles in the dynamic control of mRNA localization. We characterize the continuous regulation of nuclear speckles through a phosphorylation-dephosphorylation cycle. Shifting the balance towards dephosphorylation increases mRNA retention, consistent with increased cohesion of the structures. Although dephosphorylation of SRSF proteins has been implicated in mRNA export (Twyffels et al., 2011), our data shows lower nuclear polyA RNA levels when CLK1 or DYRK3 dissolve nuclear speckles. This suggests that export continues for many mRNAs under hyperphosphorylation. Earlier models speculated that kinases localize to condensates, while phosphatase activity occurs in the surrounding cytoplasm or nucleoplasm (Söding et al., 2020). However, our results indicate that PP1 phosphatases localize to nuclear speckles and show enrichment under conditions associated with increased condensate cohesion. This suggests that both phosphorylation and dephosphorylation occur inside or in the immediate vicinity of condensates.

mRNA enrichment within speckles varies by transcript. Using an established proximity labeling and sequencing approach for RNA, we characterized the nuclear speckle transcriptome in hiPSCs. Overall, we found that transcripts enriched in nuclear speckles tend to be more retained within the nucleus. We observed effects of increased cohesion on nuclear mRNA retention in both HeLa cells and hiPSCs. However, differences between the baseline hiPSC speckle transcriptome and the Hek293T cell speckle transcriptomes published by Barutcu et al. (2022) suggest that enrichment of individual mRNAs may differ among cell types. Transcript expression levels, which vary between cell types and tissues, were anti-correlated with nuclear speckle enrichment. A predictive model excluding expression levels as a feature and based only on invariant sequence features showed limited prediction strength. Transcripts also exhibit differential localization within nuclear speckles (Paul et al., 2022), which could influence results of proximity-based labelling approaches depending on which nuclear speckle marker is used. This may further differentiate the nuclear speckle transcriptome we describe from previously reported ones.

The retention of mRNA within the nucleus under stress conditions has previously been described in yeast, which lack nuclear speckles. There, nuclear export acts as the dominant factor in nuclear retention (Saavedra et al., 1996; Zander et al., 2016). In mammalian cells, our results suggest that the material properties of nuclear speckles can be tuned to retain mRNA in the nucleus. Similar to a role proposed for stress granules, these condensates therefore sequester mRNA away from the translation machinery during stress (Kedersha & Anderson, 2002; Moon et al., 2019). While stress granules may not exclude all translational machinery (Mateju & Chao, 2022), though, we show decreased translation under conditions with higher mRNA retention. In the acute heat stress and oxidative stress conditions we examined, we found small shifts in enrichment for overall polyA RNA, but the changes we observe under dephosphorylating conditions suggest widespread effects across genes. Heat shock has also previously been reported to affect splicing, with one paper reporting both increased and decreased intron retention and another reporting splicing inhibition (Boutz et al., 2015; Shalgi et al., 2014). Thus, intron retention could contribute to increased nuclear speckle retention for specific subsets of genes, although this remains to be confirmed.

Environmental perturbations have also been shown to affect the phosphorylation of splicing factors through changes in the expression of CLK1, as well as decreased activity of CLK1 in vitro in response to heat shock (Haltenhof et al., 2020; Jakubauskiene et al., 2015; Ninomiya et al., 2011). In the case of hypoxia, another study reported the dissolution of nuclear speckles through a decrease in SRSF6 expression (de Oliveira Freitas Machado et al., 2023). Here, we find that speckles remain intact under hypoxia in hiPSCs, although the material properties of the structures shift towards a more liquid-like state in tandem with increased phosphorylation of SR-rich proteins. Our study addresses only short perturbations that result in reversible changes in phosphorylation. Further investigation is necessary to determine whether longer term changes related to the accumulation of neurodegenerative disease-related transcripts and proteins within nuclear speckles also promote mRNA retention across the transcriptome and decreased translation (Hung et al., 2018; Jain & Vale, 2017; Lester et al., 2021). Based on our results, showing a link between phosphorylation, nuclear speckle cohesion, and mRNA nuclear retention, we propose a means through which transient changes in the nuclear retention of transcripts occur in response to stress and other environmental perturbations in mammalian cells.

## Materials and Methods

### Cell culture

HeLa-FlpIn-Trex cells were a kind gift from Ivan Dikic (Goethe University, Frankfurt) and HEK293T cells were from ATCC (Molsheim Cedex). HeLa and HEK cells were maintained in DMEM supplemented with 10% FBS and 5% L-glutamine.

hiPSCs were obtained from the Allen Cell Collection of the Coriell Institute for Medical Research (GM25256, AICS-0094-024). hiPSCs were maintained according to the standard operating procedures from the Allen Cell Institute (SOP: WTC culture v1.7) with minor adaptations. In short, hiPSCs were maintained in mTeSR Plus (STEMCELL Technologies, 100-0276). For passaging, hiPSCs were washed with DPBS and treated with Accutase (Thermo Fisher, A1110501) until they detached. Accutase treatment was quenched by dilution in DPBS. Cells were seeded on Geltrex (Thermo Fisher, A1413301) coated dishes and the growth media was supplemented with 10 µM Y-27632 (Hello Bio, HB2297) for 24 hours after replating.

Cells were grown in a humidified incubator at 37°C under 5% CO2 unless otherwise noted.

### Immunofluorescence

Cells were fixed with 4% PFA for 15-30 minutes and permeabilized with 0.5% Triton-X for 30 minutes at room temperature. Cells were then blocked for 1h at room temperature and stained with primary antibody for 1h (HeLa cells) or 2h (hiPSCs), washed in PBS, and stained with secondary antibody for 1h (HeLa cells) or 2h (hiPSCs).

#### Antibodies

1H4/pSR (Merck MABE50), B23/NPM1 (Sigma B0556), G3BP (abcam ab56574), HIF-1α (abcam ab179483), PPP1CA (abcam ab137512), SC35 (Sigma S4045), SON (ab121759).

### Plasmids

GFP-PP1-NIPP1 and GFP-PP1m-NIPP1 plasmids were a kind gift from the Bollen Lab (Bollen et al., 2010). The cargo-binding domain of TNPO3 (CBDT) plasmid was a kind gift from the Shav-Tal Lab (Hochberg-Laufer et al., 2019). GFP-phosphatase plasmids were a kind gift from the Bodenmiller Lab (Lun et al., 2019). pCMV-hyPBase was a kind gift from the Wellcome Trust Sanger Institute (Yusa et al., 2011). pECFP(C1)-NIPP1 (Addgene plasmid # 44226; http://n2t.net/addgene:44226; RRID:Addgene_44226), pEGFP(C1)-PP1alpha (Addgene plasmid # 44224; http://n2t.net/addgene:44224; RRID:Addgene_44224), pEGFP(N3)-PP1beta (Addgene plasmid # 44223; http://n2t.net/addgene:44223; RRID:Addgene_44223), and pEYFP(C1)-PP1gamma (Addgene plasmid # 44230; http://n2t.net/addgene:44230; RRID:Addgene_44230) were gifts from Angus Lamond & Laura Trinkle-Mulcahy. AICSDP-82:SON-mEGFP was a gift from Allen Institute for Cell Science (Addgene plasmid # 133964; http://n2t.net/addgene:133964; RRID:Addgene_133964). AICSDP-42:AAVS1-mTagRFPT-CAAX was a gift from Allen Institute for Cell Science (Addgene plasmid # 107580; http://n2t.net/addgene:107580; RRID:Addgene_107580). APEX2-NLS. APEX2-NLS was a gift from Alice Ting (Addgene plasmid # 124617; http://n2t.net/addgene:124617; RRID:Addgene_124617) (Kaewsapsak et al., 2017). XLone-GFP was a gift from Xiaojun Lian (Addgene plasmid # 96930; http://n2t.net/addgene:96930; RRID:Addgene_96930) (Randolph et al., 2017).

### Fluorescence recovery after photobleaching (FRAP) experiments

All FRAP experiments were performed on a Leica SP8 Falcon microscope using a 63x 1.3 NA, glycerol, Plan-Apochromat objective. SRRM2-mCh was transiently overexpressed in HeLa TREx cells in combination with various GFP- or YFP-tagged plasmids. For FRAP during drug treatments and environmental perturbations, hiPSCs were endogenously tagged with 2xGFP at the SON locus as described below. Photobleaching was performed during 50-70 minutes of DRB treatment, 90-110 minutes of GSK626616 treatment, 70-85 minutes of Pladienolide-B treatment, or after 40-85 minutes of arsenite (500 µM) treatment or 50-105 minutes of heat shock (43°C).

### Proteomics and phosphoproteomics

Hek293T cells were seeded in 10-cm dishes to reach 70% confluency at time of transfection. Triplicate technical replicates per condition per condition. Cells were transfected with 5 µg DNA (EGFP-C1, EGFP-DYRK3, GFP-PP1-NIPP1, or GFP-PP1m-NIPP1) using GeneJuice. 24h post-transfection, cells were washed twice in PBS and lysed in 450 µl lysis buffer (25 mM Tris HCl pH 7.4, 125 mM NaCl, 1 mM MgCl2, 1 mM EGTA, 5% glycerol, 1% Triton-X, 2x protease inhibitor cocktail, and 2x phosphatase inhibitor cocktail in milliQ-H2O) for 30 minutes on ice, harvested by scraping, then centrifuged at 17,000g for 10 minutes at 4°C. 25 µl anti-GFP magnetic agarose beads (Chromotek) per sample were equilibrated by washing 3x with 500 µl lysis buffer. Supernatants from the cellular lysates were then added to the beads and rotated at 4°C for 1h. Beads were washed 2x with 500 µl lysis buffer, and once with 125 µl wash buffer (1 mM Tris HCl, 125 mM NaCl, 1 mM MgCl2 in milliQ H2O).

For each sample, the anti-GFP beads with 100 µl of 10 mM Tris/2 mM CaCl2, pH 8.2 and re-suspended in 45 µl digestion buffer (triethylammonium bicarbonate (TEAB), pH 8.2), reduced with 5 mM TCEP (tris(2-carboxyethyl) phosphine) and alkylated with 15 mM chloroacetamide. Proteins were on-bead digested using 500 ng Sequencing Grade Trypsin (Promega). The digestion was carried out at 37°C overnight. The supernatants were transferred to new tubes and the beads were washed with 150 µl trifluoroacetic acid (TFA) buffer (0.1% TFA, 50% acetonitrile) and combined with the first supernatant. For controls without phosphoenrichment, 10% of the samples were dried to completeness and re-solubilized in 20 µl of MS sample buffer (3% acetonitrile, 0.1% formic acid).

For enrichment of phosphopeptides, the residual 90% of the samples were dried almost to completeness (~5 µl). The phosphopeptide enrichment was performed using a KingFisher Flex System (Thermo Fisher Scientific) and Ti-IMAC MagBeads (ReSyn Biosciences). Beads were conditioned following the manufacturer’s instructions, consisting of 3 washes with 200 µl of binding buffer (80% acetonitrile, 0.1 M glycolic acid, 5% TFA). Each sample was dissolved in 200 µl binding buffer. The beads, wash solutions and samples were loaded into 96 well deepwell plates and transferred to the KingFisher. Phosphopeptide enrichment was carried out using the following steps: washing of the magnetic beads in binding buffer (5 min), binding of the phosphopeptides to the beads (30 min), washing the beads in wash 1-3 (binding buffer, wash buffer 1 and 2, 3 min each) and eluting peptides from the beads (50 µl 1% NH4OH, 10 min). The phosphopeptides were dried to completeness and re-solubilized with 10 µl of 3% acetonitrile, 0.1% formic acid for MS analysis.

LC-MS/MS analysis was performed on an Q Exactive mass spectrometer (Thermo Scientific) equipped with a Digital PicoView source (New Objective) and coupled to a nanoAcquity UPLC (Waters Inc.). Solvent composition at the two channels was 0.1% formic acid for channel A and 0.1% formic acid, 99.9% acetonitrile for channel B. Column temperature was 50°C. For each sample 4 μl of peptides were loaded on a commercial Symmetry C18 trap column (5 µm, 180 µm x 20 mm, Waters Inc.) connected to a BEH300 C18 column (1.7 µm, 75 µm x 150 m, Waters Inc.). The peptides were eluted at a flow rate of 300 nl/min with a gradient from 5 to 35% B in 60 min, 35 to 60% B in 5 min and the column was washed at 80% B for 10 min before equilibrating back to 5% B.

The mass spectrometer was operated in data-dependent mode (DDA) using Xcalibur, with spray voltage set to 2.5 kV and heated capillary temperature at 275 °C. Full-scan MS spectra (350−1500 m/z) were acquired at a resolution of 70’000 at 200 m/z after accumulation to a target value of 3’000’000 and a maximum injection time of 100 ms, followed by HCD (higher-energy collision dissociation) fragmentation on the ten most intense signals per cycle. Ions were isolated with a 1.2 m/z isolation window and fragmented by higher-energy collisional dissociation (HCD) using a normalized collision energy of 25 %. HCD spectra were acquired at a resolution of 35’000 or 70’000 and a maximum injection time of 125 or 250 ms for phospho-enriched and non-enriched samples, respectively. The automatic gain control (AGC) was set to 3000 ions. Charge state screening was enabled and singly and unassigned charge states were rejected. Only precursors with intensity above 25’000 or 12’000 for phospho-enriched and non-enriched samples, respectively, were selected for MS/MS. Precursor masses previously selected for MS/MS measurement were excluded from further selection for 40 s, and the exclusion window tolerance was set at 10 ppm. The samples were acquired using internal lock mass calibration on m/z 371.1010 and 445.1200.

The mass spectrometry proteomics data were handled using the local laboratory information management system (LIMS) (Türker et al., 2010).

### Proteomics analysis

The acquired raw MS data were processed by MaxQuant (version 1.6.2.3), followed by protein identification using the integrated Andromeda search engine (Cox & Mann, 2008). Spectra were searched against the Uniprot Homo sapiens reference proteome (taxonomy 9606, canonical version from 2019-07-09), concatenated to its reversed decoyed fasta database and common protein contaminants. Carbamidomethylation of cysteine was set as fixed modification, while methionine oxidation, phosphor (STY) and N-terminal protein acetylation were set as variable. Enzyme specificity was set to trypsin/P allowing a minimal peptide length of 7 amino acids and a maximum of two missed cleavages. MaxQuant Orbitrap default search settings were used. The maximum false discovery rate (FDR) was set to 0.01 for peptides and 0.05 for proteins. Label free quantification was enabled and a 2-minute window for match between runs was applied. In the MaxQuant experimental design template, each file is kept separate in the experimental design to obtain individual quantitative values.

Data was then processed using R (v3.6.3). Results were first filtered to exclude reverse database hits, potential contaminants, and proteins identified only by site. Protein groups were then filtered for entries for ≥ 2 replicates in any condition under comparison. Missing LFQ intensities were imputed with random noise simulating the detection limit of the mass spectrometer (a log normal distribution with 0.25x the standard deviation of the measured, logarithmized values, down-shifted by 1.8 standard deviations). Sample differences were then tested with the t.test function in R.

For analysis of differential phosphorylation from phosphoproteomics data, significance was calculated for single sites (phosphopeptides and matched unphosphorylated peptides from input samples) by comparing generalized linear models with and without an interaction term for phosphorylation status and condition using a likelihood ratio test.

### 5-EU and pulse chase

Induction of R-MCD (+0.2)-GFP and inducible GFP hiPSCs was started 24 hours prior to the start of the experiment by supplementing the growth medium with 2 µg/ml doxycycline. Then, the cells were pulsed with fresh growth medium containing 20 µM CX5461 (MCE, HY-13323) and 1 mM 5-EU for 30 minutes. After the pulse, the cells were washed twice with growth medium and then kept in growth medium containing 1 mM Uridine for up to four hours. A new batch of cells was pulsed every hour and all cells were fixed at the same timepoint.

For 5-EU measurements under environmental perturbations, cells were similarly treated with 20 µM CX5461 and 1 mM 5-EU for 30 minutes, then fixed.

CLICK reactions to detect 5-EU were performed after fixation and permeabilization by washing cells twice in TBS, then adding a solution of 2 mM CuSO_4_, 10 µM AlexaFluor 647 Azide Triethylammonium Salt (ThermoFisher A10277), and freshly added 100 mM sodium ascorbate in TBS, and incubating in the dark at room temperature for 30 minutes before washing in PBS and proceeding with additional stainings.

### PolyA FISH

Cells were fixed and permeabilized as for immunofluorescence imaging. Cells were then washed twice with FISH wash buffer (10% formamide, 2x saline-sodium citrate (SSC)), then incubated 1h at 37°C in pre-hybridization buffer (100 mg/ml dextran sulfate, 7.5% formamide, 1.5x SSC). Half the volume was then aspirated and 2x pre-warmed hybridization buffer (100 mg/ml dextran sulfate, 10% formamide, 2x SSC, 800 nM dT-30-Atto488, dT-30-Cy3, or dT-30-Cy5 oligomer) added in equal volume for overnight incubation at 37°C. The following day, cells were washed twice for 30 minutes at 37°C with pre-warmed FISH wash buffer, then washed with 2x SSC and finally PBS before proceeding with immunofluorescence staining and imaging.

### Design of smFISH probes

Probes for smFISH were designed as branched DNA probes for signal amplification. Twelve primary probes were designed per target gene using PaintSHOP (Hershberg et al., 2021), using the hg38 newBalance (isoform flattened) probe set with default settings or OligoMiner (Beliveau et al., 2018; Passaro et al., 2020). When more than twelve probes were found for a target, the probes with highest on_target value (PaintSHOP) or lowest melting temperatures (OligoMiner) were chosen. For three gene targets, fewer than 12 probes were available (EXOSC1: 8, H2BC11: 8, POU5F1: 6). For pooled probes, around 50 gene targets were selected per pool (enriched pool: 48, depleted pool: 47, neither pool: 48) based on relaxed thresholds of significance (enriched and depleted pools: |log2 fold change| > 0.3, padj < 0.6, neither pool: |log2 fold change| < 0.05, padj > 0.99). The primary probes were then extended by color-specific barcodes to which four secondary probes could bind. Primer binding sequences for probe purification were also included. The signal was amplified in this manner up to quaternary probes. Finally, fluorophore labelled probes (label probes) were bound to quaternary probes. Color specific barcodes were derived from orthogonal 25mer barcode sequences designed previously (Xu et al., 2009). The labelled probes were based on sequences used in the amplification method for smFISH signals described previously (Kishi et al., 2019). Raw probe sequences as generated by PaintSHOP and OligoMiner as well as sequences extended with barcodes are listed in Supplementary Table 2. Probe sequences for amplification probes (secondary, tertiary, quaternary and label probes) are listed in Supplementary Table 4. Primary probes were ordered from Twist Bioscience or Integrated DNA Technologies (IDT) as oligo pools. Amplification probes were ordered from Microsynth.

### Purification of oligo pools

FISH probes were purified from oligo pools in three steps. First, oligo pools were amplified by PCR using primers to introduce T7 RNA polymerase promoter sequences and the amplicon was cleaned up using Zymo DNA Clean & Concentrator-25 (Zymo Research, D4033). The resulting amplicon was used as starting material for in vitro transcription using HiScribe T7 High Yield RNA Synthesis Kit (NEB, E2040S). Finally, the RNA product was used for reverse transcription using Maxima H Minus Reverse Transcriptase (Thermo Fisher, #EP0752) and RNA was removed by alkaline lysis.

### smFISH

hiPSCs were grown on 96-well plates (Greiner, 655090) and fixed with 4% PFA (EMS, 15710) in PBS for 15 minutes at room temperature. When pooled probe sets were used, to ensure even entry of probes, cells were dissociated and seeded as single cells before the experiment. The sample was then washed with PBS and permeabilized with 0.5% Triton X-100 (Sigma Aldrich, X100) in PBS for 30 minutes at room temperature. After another PBS wash, if pooled probe sets were used, the sample was treated with Protease QS (Thermo Fisher, QVP0011) diluted 1:2000 in PBS for 10 minutes at room temperature while shaking. Protease treatment was stopped by washing the sample with protease stop buffer (Thermo Fisher, QVP0011) twice and once with PBS. The sample was then incubated with 40% wash buffer (40% Formamide, 2x SSC, 0.001% tween20 in RNase free H_2_O) for one hour at 65°C. The sample was then incubated with primary probes diluted to a final concentration of 2 nM per probe in primary probe hybridization solution (10% Dextran, 40% Formamide, 2x SSC, 0.01% yeast tRNA, murine RNase inhibitor, 0.001% tween20 in RNase free H_2_O) for 16 hours at 37°C. The sample was washed three times with 40% wash buffer and once with 30% wash buffer (30% Formamide, 2x SSC, 0.001% tween20 in RNase free H_2_O) for 6 minutes at 37°C. The sample was then incubated with secondary probes diluted to 5 nM in probe solution (10% Dextran, 40% Formamide, 2x SSC, 0.001% tween20 in RNase free H_2_O) for 30 minutes at 37°C. After incubation with secondary probes, the sample was washed three times with 30% wash buffer for 6 minutes at 37°C. Probe addition and washing were repeated in the same manner for tertiary and quaternary branching probes. The sample was then washed with PBS and incubated with label probes diluted to 0.5 µM in PBS for 1 hour at 37°C protected from light.

### Nucleofection of hiPSCs

Y-27632 was added to hiPSC growth media 1-6 hours prior to nucleofection at a final concentration of 10 µM. Cells were then passaged as previously described and 8×10^5^ cells were resuspended in 100 µl of nucleofector solution (Lonza, VPH-5012). Depending on the experiment, plasmids, sgRNA and Cas9 were added to the resuspended cells. The cell suspension was then transferred to a nucleofection cuvette and nucleofection was performed using Lonza Nucleofector 2b using program A-023. After nucleofection, 500 µl of warm growth medium was added to the nucleofection cuvette and cells were transferred to a Geltrex-coated well of a 6-well plate containing pre-equilibrated (37°C, 5% CO_2_) mTeSR plus supplemented with 10 µM Y-27632. Y-27632 concentration in the growth medium was halved to 5 µM after 24 hours, and it was removed completely after 48 hours of plating.

### Design of sgRNA and Donor Plasmids for APEX2-SON and APEX2-NLS hiPSCs

We used crRNA sequences specified by the Allen Cell Institute to guide Cas9 to the SON locus and the AAVS1 locus. We ordered sgRNAs incorporating the crRNA and tracrRNA sequences from Sigma Aldrich. We used plasmid AICSDP-82:SON-mEGFP as template for the APEX2-SON donor plasmid and AICSDP-42:AAVS1-mTagRFPT-CAAX as a template for the APEX2-NLS donor plasmid. These plasmids already contained the homology arms needed for downstream CRISPR/Cas9 genome editing. AICSDP-42:AAVS1-mTagRFPT-CAAX additionally contained the CAG promoter to allow stable and consistent transgene expression in hiPSCs (Luo et al., 2014; Oceguera-Yanez et al., 2016). The APEX2 enzyme sequence and NLS motif were PCR amplified from plasmid APEX2-NLS. The APEX2 sequence was inserted in front of mEGFP in AICSDP-82:SON-mEGFP to create the APEX2-mEGFP-SON donor plasmid. The APEX2-NLS sequence was inserted along with the mEGFP sequence from AICSDP-82:SON-mEGFP into AICSDP-42:AAVS1-mTagRFPT-CAAX, replacing mTagRFPT-CAAX, to create the mEGFP-APEX2-NLS donor plasmid. All donor plasmids were assembled by Gibson assembly using NEB Gibson Assembly Master Mix (NEB, E2611L).

### crRNA sequences

SON: CTGCTCGATGTTGGTCGCCA

AAVS1: GGGGCCACTAGGGACAGGAT

### Cell line generation

The protocols for CRISPR/Cas9 genome engineering in hiPSCs were adapted from published protocols (Haupt et al., 2018; Roberts et al., 2017). To generate the 2xGFP-SON, APEX2-NLS, and APEX-SON cell lines, hiPSCs were nucleofected as described above with 2 µg of donor plasmid and 1.5 µl each of 10 µM sgRNA and 10 µM Cas9 (Sigma Aldrich, CAS9PROT-50UG). Once cells reached confluency, they were passaged, resuspended in phenol red free mTeSR1 (STEMCELL, 05876) and sorted by FACS for GFP positive cells.

To generate doxycycline inducible R-MCD (+0.2)-GFP and GFP cell lines, hiPSCs were nucleofected as described above with 1 µg of XLone-R-MCD (+0.2)-GFP or XLone-GFP plasmid along with 1 µg of PiggyBack transposase plasmid (pCMV-hyPBase). XLone-R-MCD (+0.2)-GFP plasmid was generated by PCR amplification of R-MCD (+0.2) domain from pmEGFP-N1-R-MCD(+0.2) plasmid (kind gift from Dr. Gregory Jedd) and cloned into XLone-GFP plasmid using Gibson assembly kit (NEB, E2611S). Two days after nucleofection, selection was started by adding blasticidin (Santa Cruz, sc-495389) at a concentration of 10 μg/ml. Blasticidin selection was carried out for multiple passages before cells were frozen in liquid nitrogen for long-term storage.

### Drug treatment

Drug treatment with GSK626616 (5 µM), DRB (75 µM) and Pladienolide-B (2 µM) was carried out in DMEM/F-12 medium without serum for two hours at 37°C and 5% CO_2_. For DYRK3 and CLK1 inhibition in hypoxia, GSK626616 (5 µM), TG003 (100 µM), or DMSO were likewise diluted in serum-free medium (pre-equilibrated overnight under hypoxic conditions) and added to cells for two hours after 22 hours in hypoxia (0.2% O2, 5% CO2, 37°C). Inhibition with CLK-IN-T3 (1 µM) was carried out in complete medium for 8 hours (OPP experiment) or 12 (FISH experiments) hours.

### OPP

Translation was quantified based on the incorporation of O-propargyl puromycin (OPP) in nascent peptides (Liu et al., 2012). Cells were incubated in media containing 30 µM OPP (with drugs) for 45 minutes before fixation and detection of OPP through a CLICK reaction, as described above for 5-EU.

### Stresses and environmental perturbations

Heat shock was performed at 43°C for 1 hour in a cell culture incubator maintained at 5% CO2. Oxidative stress was induced for 1 hour using 500 µM sodium meta-arsenite dissolved in media. To induce hypoxia, cell media was exchanged for media pre-equilibrated under hypoxic conditions (0.2% O2), and cells were maintained at 0.2% O2/5% CO2 in a humidified atmosphere at 37°C in a hypoxia workstation (Baker-Ruskinn).

### APEX2 enzymatic reaction

The protocol for the APEX2 enzymatic reaction was adapted from Fazal et al., 2019 and Padrón et al., 2019. APEX2-tagged hiPSCs were washed with DPBS (Thermo Fisher, 14190136) and then incubated in DPBS containing 0.5 mM biotin-phenol (Iris Biotech, LS-3500.1000) and 0.005% digitonin (Sigma Aldrich, D141) for 3 minutes at room temperature. To trigger the enzymatic reaction, hydrogen peroxide (Sigma Aldrich, 1.07209) was added to a final concentration of 0.5 mM and the dish was tilted for 1 minute at room temperature. To stop the reaction, cells were washed once with quenching solution (5 mM Trolox, 10 mM sodium ascorbate, 10 mM sodium azide in DPBS) and three times with wash solution (5 mM Trolox, 10 mM sodium ascorbate in DPBS).

### RNA extraction and enrichment of biotinylated RNA

The protocol for extraction and enrichment of biotinylated RNA was adapted from Fazal et al., 2019. hiPSCs were lysed by adding RNA lysis buffer (Zymo Research, R1060-1-50) directly to the cell culture dish. The cells were scraped into solution and total RNA was purified using Zymo Quick-RNA Miniprep kit (Zymo Research, R1054). Of the isolated total RNA, 5 µl was set aside as an input sample. To purify biotinylated RNA, we used Pierce Streptavidin Magnetic Beads (Thermo Fisher, 88816). First, 30 µl of beads per sample were resuspended in Binding and Washing (B&W) buffer (5 mM Tris-HCL (pH 7.5), 1 mM EDTA, 2 M NaCl) by vortexing, and then washed three times with B&W buffer. The beads were then resuspended in Solution A (0.1 M NaOH, 0.05 M NaCl) and incubated for 2 minutes. Then the beads were washed twice with Solution B (0.1 M NaCl) and resuspended in 100 µl Solution B. An equal volume of total RNA sample was added, and the samples were incubated for 2 hours at 4°C on a rotator to allow the biotinylated RNA to bind to the beads. The beads were then washed three times with B&W buffer and resuspended in 54 µl of RNase free H_2_O. A 3X proteinase buffer was prepared (For 1 ml: 300 µl PBS, 300 µl 20% N-Lauryl sarcosine sodium solution (Sigma Aldrich, L7414), 60 µl 0.5 M EDTA, 15 µl 1 M DTT, 225 µl RNase free H_2_O), and 33 µl of this buffer was added to the beads together with 10 µl Recombinant Proteinase K Solution (20 mg/ml, Thermo Fisher, AM2548) and 3 µl RiboLock RNase inhibitor (Thermo Fisher, EO0381). The beads were then incubated for 1 hour at 42°C and then for 1 hour at 55°C on a shaker. Finally, biotinylated RNA was purified using RNA Clean & Concentrator-5 kit (Zymo Research, R1013).

For biotinylated RNA, five biological replicates were collected per cell line and drug condition. For input RNA, three biological replicates were collected per cell line and drug condition.

### Subcellular fractionation of hiPSCs

The protocol used for subcellular fractionation of hiPSCs was adapted from Mayer & Churchman, 2017. Throughout the protocol, samples were kept at 4°C and handled under RNase free conditions. Briefly, hiPSCs were grown to confluency on 10 cm dishes and were first washed with and then scraped into DPBS. Cells were then pelleted by centrifugation for 3 minutes at 211 g and resuspended in lysis buffer (0.15% NP-40, 150 mM NaCl, 25 µM α-amanitin, 10 U SUPERase.IN (Thermo Fisher, AM2696), 1x cOmplete protease inhibitor mix, EDTA free (Sigma Aldrich, 11873580001)). The lysate was layered onto a sucrose buffer (10 mM Tris-HCl (pH 7.0), 5 M NaCl, 25% (w/v) sucrose, 25 µM α-amanitin, 10 U SUPERase.IN, 1x cOmplete protease inhibitor mix, EDTA free) and centrifugated for 10 minutes at 16000 g. The supernatant representing the cytoplasmic fraction was collected and the remaining pellet was washed twice with nuclei wash buffer (1 mM EDTA, 25 µM α-amanitin, 40 U SUPERase.IN, 1x cOmplete protease inhibitor mix, EDTA free prepared in PBS). The nuclear pellet was then resuspended in glycerol buffer (20 mM Tris-HCl (pH 8.0), 75 mM NaCl, 0.5mM EDTA, 50% (v/v) glycerol, 0.85 mM DTT, 25 µM α-amanitin, 10 U SUPERase.IN, 1x cOmplete protease inhibitor mix, EDTA free) to which nuclei lysis buffer (1% NP-40, 20 mM HEPES (pH 7.5), 300 mM NaCl, 1 M urea, 0.2 mM EDTA, 1 mM DTT, 25 µM α-amanitin, 10 U SUPERase.IN, 1x cOmplete protease inhibitor mix, EDTA free) was added. After 5 minutes of incubation the suspension was centrifugated at 18500 g for 2 minutes. The supernatant representing the nuclear fraction was collected and the remaining pellet was washed with PBS and centrifugated for 1 minute at 1150 g. The supernatant was discarded, and the pellet representing the chromatin fraction was resuspended in 50 µl of PBS. The chromatin fraction was then incubated with TRIzol (Life Technologies, 15596) and chloroform for 5 minutes at room temperature. The sample was centrifuged, and the upper aqueous phase was collected.

To isolate RNA, 3.5 sample volumes of RLT buffer (Qiagen, 79216) were added to chromatin, nuclei and cytoplasm fractions. After mixing, 2.5 volumes of ice-cold 75% ethanol was added. RNA was then cleaned up using the RNeasy Mini kit (Qiagen, 74104). A total of 5 biological replicates were collected.

### Library preparation and RNA sequencing

For APEX2-sequencing samples, libraries were prepared using SMARTer Stranded Total RNA-Seq Kit v2 - Pico Input Mammalian (Takara, 634411). Single-end sequencing was performed on the Ilumina NovaSeq platform with a sequencing depth of 80 million reads and a read length of 100 bp.

For subcellular fraction sequencing samples, libraries were prepared using Truseq Stranded mRNA kit (Illumina, 20020594). Single-end sequencing was performed on the Illumina NovaSeq platform with a sequencing depth of 200 million reads and a read length of 100 bp. Library preparation and RNA sequencing was performed by the Functional Genomics Center Zurich (FGCZ).

### Processing of RNAseq data

Trimmomatic v0.39 (Bolger et al., 2014) was used to trim adapter sequences from raw reads with the following settings: ILLUMINACLIP:TruSeq3-SE:2:30:10 LEADING:3 TRAILING:3 SLIDINGWINDOW:4:15 MINLEN:36. Quality checks were carried out before and after trimming with FastQC v0.11.9. Trimmed reads were mapped to the human reference genome (hg38, GRCh38.p14 primary genome assembly) using GENCODE v40 gene annotations with STAR v2.7.3a (Dobin et al., 2013). Count tables were generated using the featureCounts (Liao et al., 2014) function of the R package Rsubread v2.10.5 (Liao et al., 2019).

### Differential Gene Expression analysis

Genes with less than 10 counts in any of the samples were removed prior to the analysis. Differential Gene Expression (DGE) analysis was performed with DESeq2 v1.36.0 (Love et al., 2014) using the default Wald test. To quantify enrichment in nuclear speckles vs. nucleoplasm in DMSO treatment, we tested for the influence of the cell line factor. Thresholds of |log2FoldChange| > 0.5 and padj < 0.05 were used. To quantify enrichment in nuclear speckles vs. Nucleoplasm in drug treated vs. DMSO treated cells we tested for the influence of the interaction term treatment:cell line. Thresholds of |log2FoldChange| > 0.5 and padj < 0.1 were used. Results of DESeq2 analyses are listed in Supplementary Table 1.

### Differential Transcript Expression analysis

Transcript quantification was performed using Salmon v0.12.0 (Patro et al., 2017) with the -- numGibbsSamples option set to 30 to generate Gibbs samples. Differential transcript expression analysis was performed using Fishpond v2.2.0 (Zhu et al., 2019).

### Normalization with input samples

Log2FoldChanges for input sample nuclear speckle enrichment was quantified with DESeq2 as described above. DGE results for nuclear speckle enrichment were normalized by fitting a linear regression model with log2FoldChanges of input samples as independent variable and log2FoldChanges of biotinylated samples as dependent variable. The residuals of the linear model were used as corrected log2FoldChange.

### Gene Set Enrichment Analysis

Gene Set Enrichment analysis was performed with the GSEA software v4.2.3 (Mootha et al., 2003; Subramanian et al., 2005). GSEA was run in pre-ranked mode with default settings and using KEGG, Reactome and GO:CC, GO:BP and GO:MF gene sets. As ranking metric, we used input normalized log2FoldChange in case of nuclear speckle enrichment in DMSO condition and −log_10_(padj) * sign(log2FoldChange) in case of nuclear speckle enrichment in drug vs. DMSO conditions.

### Splicing analysis

Differences in splicing between the transcripts enriched in nuclear speckles or the nucleoplasm were assessed in DMSO control samples using three tools: iREAD (Li et al., 2020), VAST v2.5.1 (Han et al., 2017; Tapial et al., 2017), and MAJIQ v 2.4.dev3+g85d0781.d20220721 (Vaquero-Garcia et al., 2016).

For VAST, analysis was run starting with untrimmed reads, as recommended, with thresholds set for detection in ≥1 samples with ≥ 10 reads, with minimum probability ≥ 0.95. Residual differences in retention were calculated using input samples from each cell line and a residual difference of > 0.15 used for plotting. Control introns (N = 10,000) were selected at random from all detected introns for feature comparisons.

For MAJIQ, detection thresholds were set for observation of alternative splicing in ≥ 1 sequencing replicate with ≥ 5 reads per junction (prior-minreads) and ≥ 2 reads per experiment (minreads), along with the default probability threshold for local splice variation of 0.2. Control introns (N = 3,633) were introns with absolute differential percent spliced in index < 0.1 and probability < 0.2. For comparison between drug-treated cells and controls (**Supplementary Figure 5G**), splicing differences were calculated using MAJIQ for SON-enriched samples under drug treatments compared to DMSO.

### Network visualization of GSEA results

Network visualizations of GSEA results were made using Cytoscape (Shannon et al., 2003) v3.9.1 and EnrichmentMap (Merico et al., 2010). Only Nodes with gene set sizes between 29 and 496 and NES smaller than −2 or greater than 1.8 are displayed. Annotations were generated using the AutoAnnotate plugin (Kucera et al., 2016) and manually adapted. Annotated groups were positioned manually.

### Prediction of transcript-level nuclear speckle enrichment

Feature categories used for the linear model were collected from the following sources. General sequence features: custom Python script using GENCODE v43 primary assembly gff3 annotations. As part of the general sequence features, we also included the GGACU m6A motif density and the AGCCC nuclear localization motif (B. Zhang et al., 2014) density. RNA binding protein features: oRNAment database (Benoit Bouvrette et al., 2020). Kinetic rates: (Smalec et al., 2022). Promoter motifs: The Eukaryotic Promoter Database EPD (Périer et al., 1998). TPM: Salmon quantification (**Supplementary Table 3**).

Features for training were filtered to only include protein coding transcripts. Features representing motif counts were transformed into density. Features that contained 0 values for more than 75% of the data were removed. All features were log transformed, missing values were imputed by the mean of the feature and features were z-scored. Principal Component Analysis (PCA) (Maćkiewicz & Ratajczak, 1993) was performed on the resulting feature set and the number of principal components explaining 99% of the variance were retained (n=414). All transcripts annotated to the same gene were only allowed to be in either the test or the train set. The reported coefficients of determination were calculated by averaging results of 10-fold cross-validation. Preprocessing and model training were done in Python (v3.9.7) using scikit-learn (v1.3.0).

### Feature importance scoring

The linear regression model was trained for one hundred iterations with different test (15% of the data) and train (85% of the data) subsets. The top ten loadings of the ten principal components with the highest absolute coefficients were extracted for each iteration. The frequency at which a feature occurs in this subset of features was taken as a measure of feature importance. For example, a feature occurring in the top 10 loadings of every single top 10 PC will have a frequency of occurrence of 1.

### UMAP representation

Transcript features were preprocessed as described above and were used as the input for the UMAP algorithm as implemented in Python (McInnes, Healy, & Melville, 2018; McInnes, Healy, Saul, et al., 2018).

### Immunofluorescence and FISH Microscopy

Microscopy images were acquired on a CellVoyager 7000 microscope (Yokogawa) equipped with an enhanced CSU-W1 spinning disk (Microlens-enhanced dual Nipkow disk confocal scanner, wide view type) and Andor cSMOS cameras or on a CellVoyager 8000 microscope (Yokogawa) equipped with CSU-W1 spinning disk and ORCA-Flash4.0 V3 cameras. Images were acquired using a 60x Nikon water immersion objective (NA=1.2) or 40x Nikon air objective (NA=0.95) on the CellVoyager 7000, and using a UPLSAP60xW customized Yokogawa objective (NA=1.2) on the CellVoyager 8000. Z-stacks with a 1 μm spacing were acquired per site, spanning the whole height of the sample (12 – 20 μm). For subsequent processing, maximum intensity projections (MIPs) were performed for each site, except where noted.

### Image Processing

Nuclear speckles were segmented based on SC35 or SON intensity images using the pixel classifier functionality of Ilastik v1.4.0 (Berg et al., 2019; Sommer et al., 2011). Nuclei were segmented based on DAPI intensity images using Cellpose (Stringer et al., 2020). Image processing was performed on a computing cluster (ScienceCloud) (https://www.zi.uzh.ch/en/teaching-and-research/science-it/computing/sciencecloud.html) provided by the Service and Support for Science IT (S3IT) facility of the University of Zurich (UZH) using TissueMAPS (https://github.com/pelkmanslab/TissueMAPS) and custom Python scripts.

### Pseudocoloring of smFISH images

Intensity values of smFISH images were mapped to different color gradients depending on the subcellular localization of the signal. Signal overlapping with nuclear speckle segmentation (based on SON-mEGFP or SC35 antibody staining) was mapped to an orange gradient, while signal not overlapping with speckles was mapped to a blue gradient. Intensities for the same smFISH target were always scaled the same way across both gradients. Color gradients were applied in Python using the microfilm package v0.2.1.

## Supporting information

Readme

## Acknowledgements

We thank all members of the Pelkmans Lab for discussions and input, particularly Cheng-Han Yang and Quentin Szabo for their contributions to the development of the smFISH protocol, and Raffaella Gallo and Yixuan Fu for DYRK3 plasmids. We also acknowledge Roland Wenger, Patrick Spielmann, and Agnieszka Jucht for assistance with hypoxia experiments. We thank the Functional Genomics Center Zurich for support in performing proteomics and sequencing experiments (in particular Catherine Aquino, Antje Dittmann, Tobias Kockmann, Laura Kunz, and Timothy Sykes), and the Center for Microscopy and Image Analysis for training for FRAP experiments and for FACS sorting. A.M. was supported by EMBO ALTF 455-2020 and is supported by MSCA individual fellowship 101029269. K.M was supported by EMBO ALTF 978-2019 and is supported by HFSP long term fellowship LT000086/2020-L. This work was supported by the Swiss National Science Foundation (grant 310030_192622) and the University of Zurich.

## Author contributions

Conceptualization, A.M., L.P., A.T.; Methodology, K.F., A.M., L.P., A.T.; Investigation, A.M., K.M., A.R., A.T., S.W.; Writing, A.M., L.P., A.T.; Visualization, A.M., A.T.; Funding Acquisition, L.P.

## Declaration of interests

L.P. consults for Dewpoint Therapeutics and has ownership interest in Sagimet Biosciences and Apricot Therapeutics.

## Supplementary Figures

**Supplementary Figure 1.**
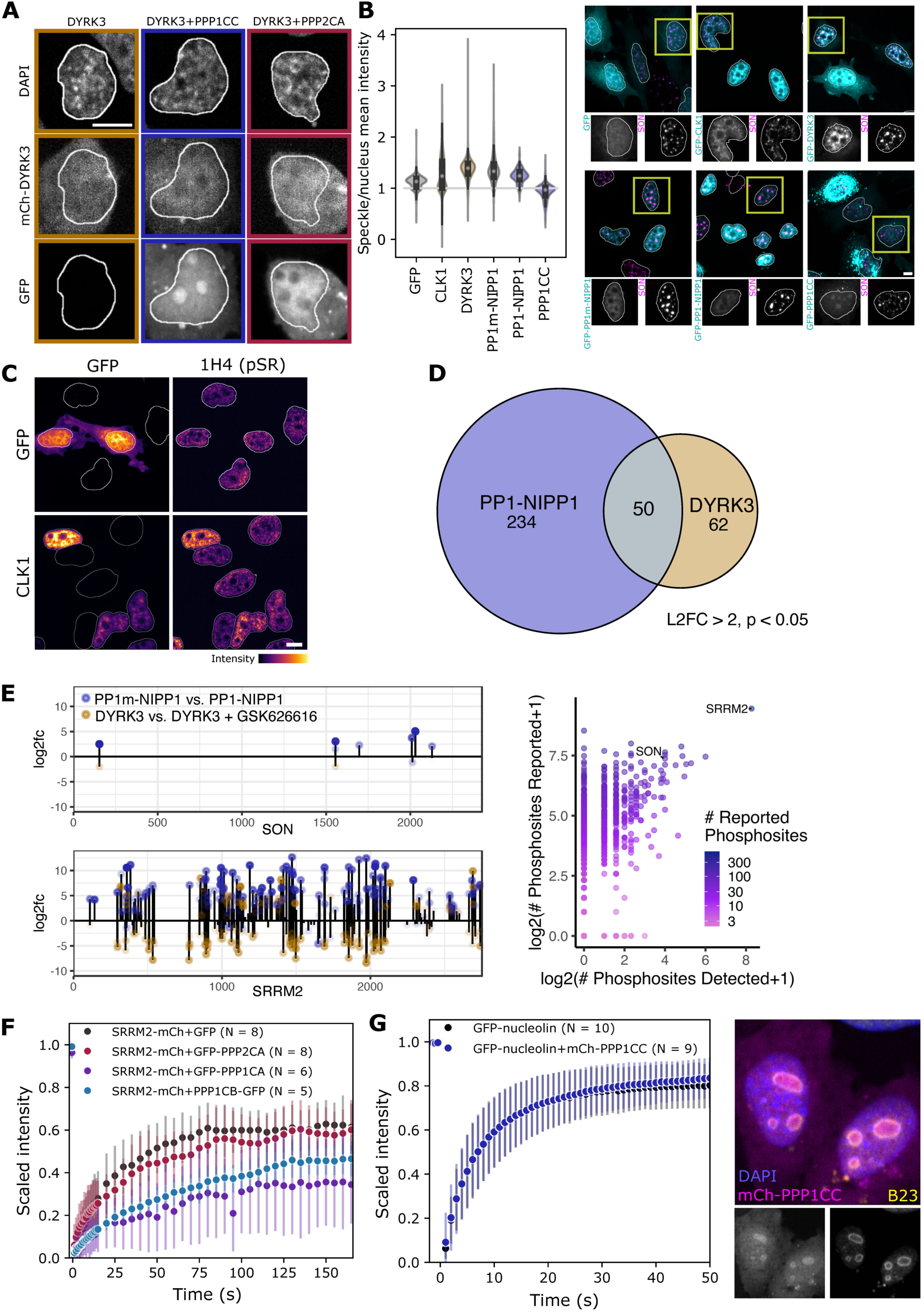
**(A)** DAPI, mCh-DYRK3, and GFP-PPP1CC imaging for example cells shown in Fig 1B either transfected with mCh-DYRK3 alone (left) or co-transfected with GFP-PPP1CC (centre) or GFP-PPP2CA (right). **(B)** (Left) Quantification of mean GFP intensity in nuclear speckles compared to the nucleus as a whole for GFP-tagged regulatory kinases and phosphatases. The regulatory subunit NIPP1 increases PPP1CC enrichment within nuclear speckles, independent of phosphatase catalytic activity. (Right) Example images, with GFP intensity scaled independently per construct. **(C)** Example images of phospho-SR staining with GFP and GFP-CLK1 overexpression. **(D)** The overlap between sites across proteins differentially phosphorylated with GFP-PP1-NIPP1 and mutant catalytic GFP-PP1m-NIPP1 expression, and sites differentially phosphorylated with GFP-DYRK3 overexpression or GFP-DYRK3 + inhibitor (GSK626616). **(E) Left:** Differentially phosphorylated sites across the scaffold proteins SRRM2 (12 co-regulated sites) and SON depending on PP1 phosphatase and DYRK3 kinase activity. **Right**: The number of phosphorylation sites we detect per protein vs. the number of reported phosphorylation sites in the database PhosphoSite Plus (Hornbeck et al., 2015). **(F)** Fluorescence recovery after photobleaching for SRRM2-mCh in HeLa cells with GFP or GFP-phosphatase overexpression shows effects on nuclear speckle cohesion by PP1 catalytic subunits but not PPP2CA. **(G)** Fluorescence recovery after photobleaching for GFP-nucleolin in HeLa cells with or without mCh-PPP1CC overexpression shows no effects of PP1 on nucleolar cohesion, despite localization of the phosphatase to nucleoli (images right of plot). Error bars for FRAP experiments show standard deviations. Scale bars correspond to 10 µM.

**Supplementary Figure 2.**
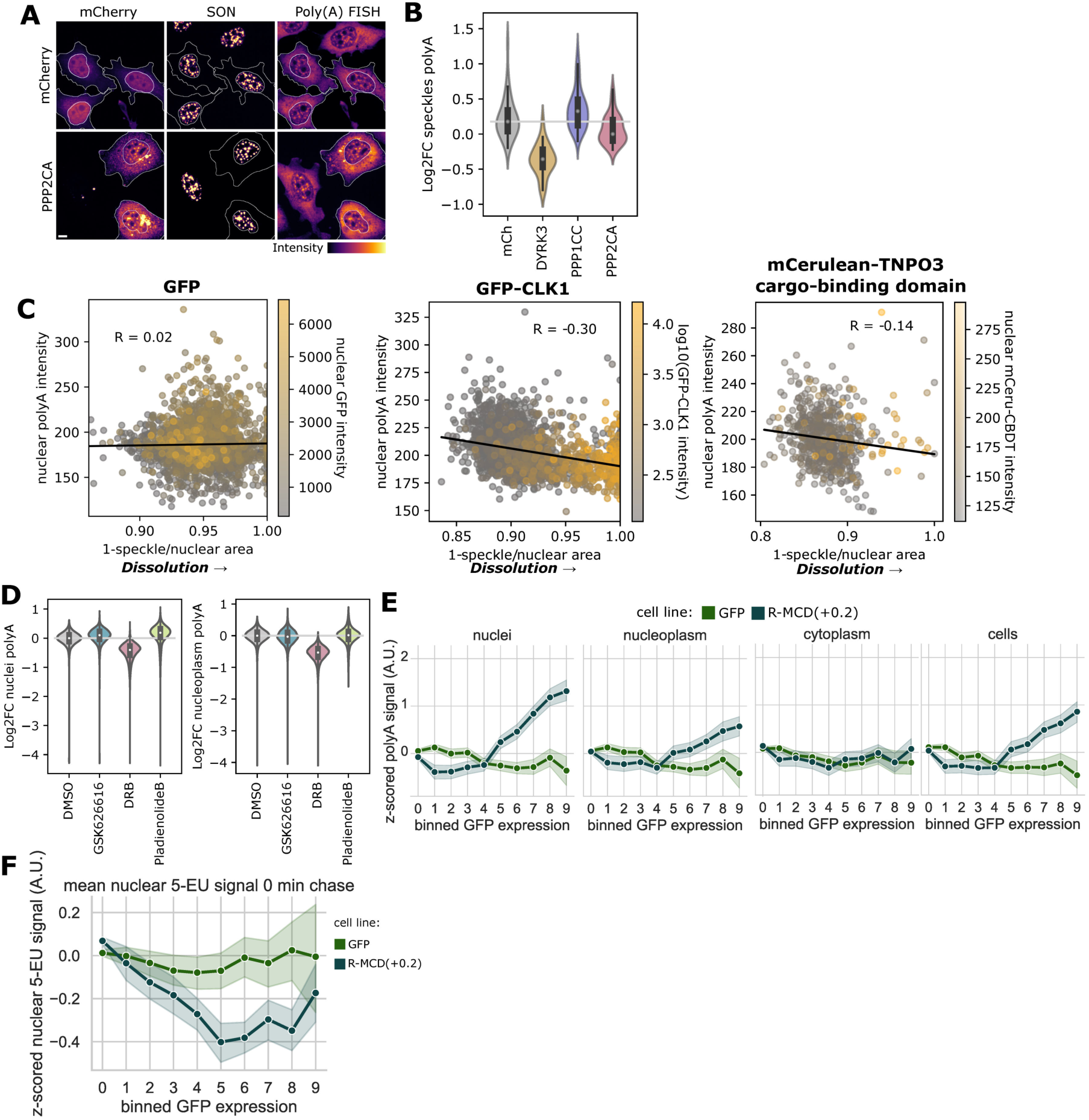
**(A)** Example images for polyA FISH signal with overexpression of mCh and mCh-PPP2CA, with outlines shown for transfected cells. **(B)** Alternative quantification of mean polyA FISH intensity across cells with high expression (top 5% of measured cells) of kinases or phosphatases within nuclear speckles showing log2 fold change compared to untransfected cells (bottom 15% of cells in mCh expression) in the same well. **(C)** Nuclear polyA intensity negatively correlates with dissolution of nuclear speckles through GFP-CLK1 and mCerulean-TNPO3 cargo-binding domain overexpression but is not affected by GFP overexpression. **(D)** Quantification of mean nuclear polyA FISH signal under DYRK3 inhibition, splicing inhibition, and transcriptional inhibition. **(E)** PolyA intensity in different subcellular localizations after 30 minutes of incubation with 5-EU and 2 hours of chase. Nuclear speckle and nucleoplasmic polyA FISH intensities are elevated depending on the expression level of a positively charged arginine-rich mixed charge domain (R-MCD+0.2), while the cytoplasmic signal intensity is not affected. **(F)** 5-EU staining with and without the expression of a positively charged mixed charge domain (R-MCD+0.2) shows that high levels of R-MCD+0.2 depress transcription.

**Supplementary Figure 3.**
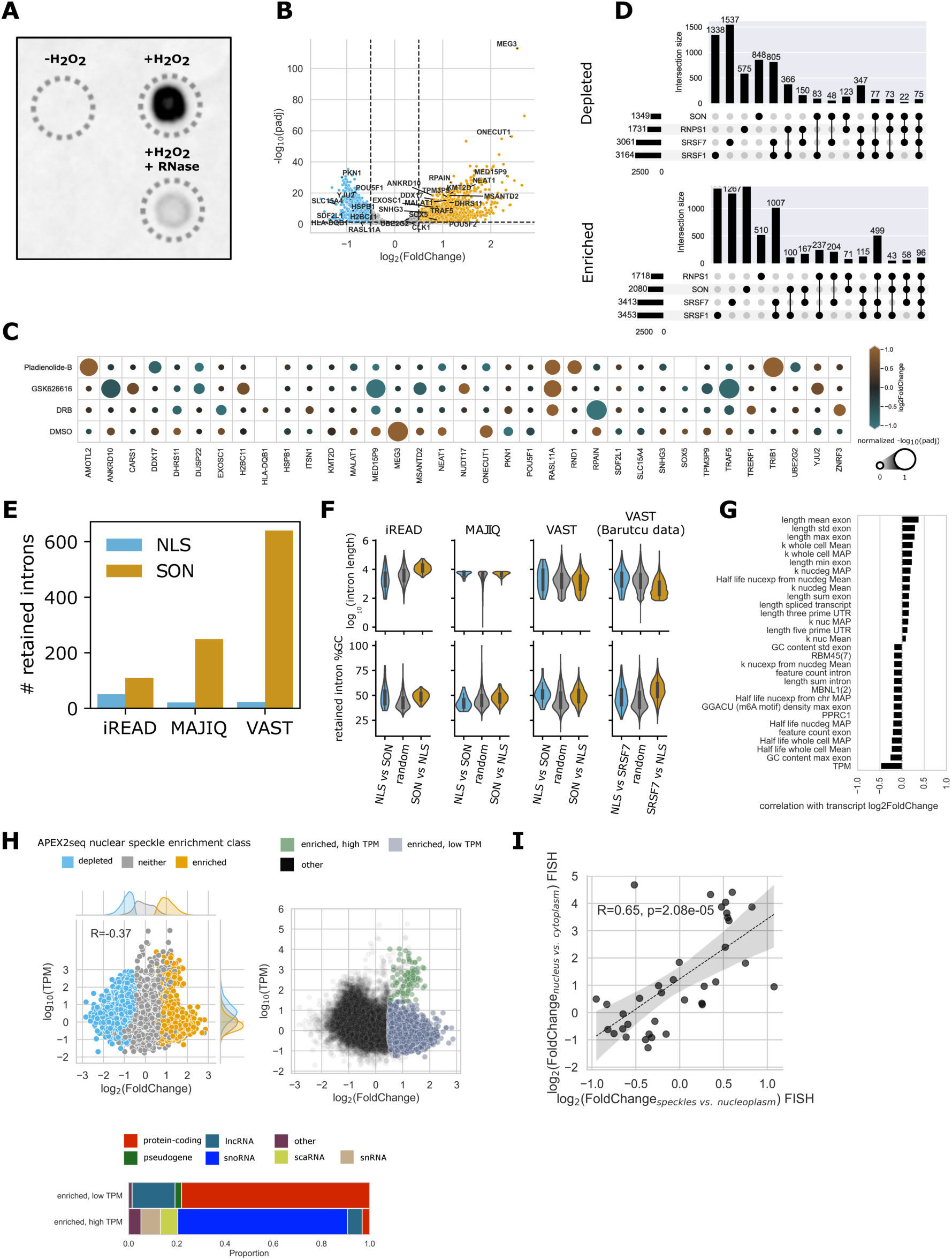
**(A)** RNA dot blot of biotin enriched RNA samples. Omitting hydrogen peroxide treatment results in no detectable biotinylation of biomolecules. RNase treatment strongly reduces the detected signal. **(B)** Volcano plot showing genes with significant nuclear speckle enrichment or depletion after input normalization (|log2FoldChange| > 0.5 and padj < 0.05). **(C)** Overview of genes against which smFISH probes were designed. Color indicates log2FoldChange of nuclear speckle enrichment as measured by APEX2-seq. Size indicates −log10(padj) normalized between 0 and 1 in each treatment condition. Rows represent gene targets and columns represent speckle enrichment under different treatment conditions (DMSO: baseline speckle enrichment, GSK626616, DRB and Pladienolide: changes in enrichment observed between treatment and DMSO condition). **(D)** Upset plots comparing hits detected as depleted (top) and enriched (bottom) with results obtained by Barutcu et al., 2022. Overlaps were small, even between nuclear speckle markers used in the same study. **(E)** More retained introns are detected as enriched within nuclear speckles (SON APEX2 data) than within the nucleoplasm (NLS APEX2 data), using three approaches to splicing analysis (iREAD, MAJIQ, and VAST). **(F)** Retained intron features show variable length differences depending on analysis methods, but consistent bias to higher GC content in nuclear speckles across detection tools and in a second dataset. **(G)** Pearson correlation coefficients of top 15 nuclear speckle enrichment correlated or anti-correlated features. **(H)** Nuclear speckle enriched transcripts mostly display lower levels of expression (TPM) than nuclear speckle depleted transcripts. A subset of nuclear speckle enriched transcripts is highly expressed. Separating this subset by fitting a gaussian mixture model shows that it mostly consists of snoRNA, scaRNA and snRNA. **(I)** Genes that show nuclear speckle enrichment over the nucleoplasm tend to be more enriched in the nucleus compared to the cytoplasm of cells.

**Supplementary Figure 4.**
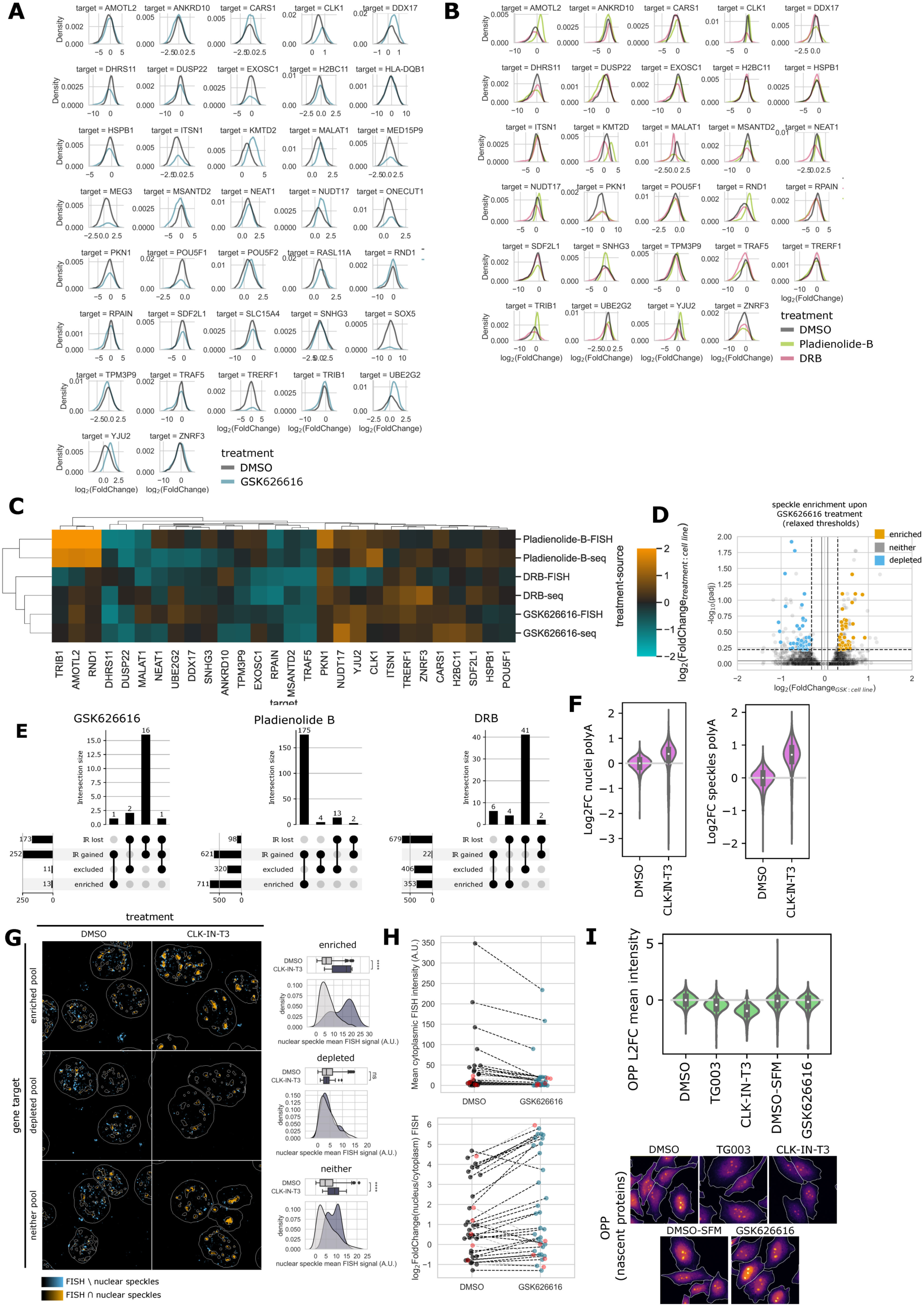
**(A)** Single cell distributions of nuclear speckle enrichment of FISH signal per target gene, colored by treatment. **(B)** Same as Supplementary Figure 4B, but for Pladienolide-B and DRB treated samples. **(C)** Clustermap of nuclear speckle enrichment under different drug treatments measured by either FISH or APEX2-seq. Results of FISH and sequencing treated with the same drug cluster together. Only gene targets that were present in all experiments are displayed. **(D)** Volcano plot visualization of the genes selected for the smFISH pooled probe sets. **(E)** UpSet plots showing the intersections between genes differentially enriched in or excluded from nuclear speckles under drug treatments, and genes with differential intron retention (IR) under drug treatments. **(F)** PolyA FISH shows retention of mRNA in nuclear speckles and the nucleus under CLK inhibition with 12h of CLK-IN-T3 treatment (1 µM). **(G)** Example pseudocolor images of FISH signal of pooled probe sets (n**≈**50 targets per pool for genes depleted, enriched, or neither upon GSK626616 treatment) in cells treated with CLK-IN-T3 for 12 hours. Density plots show single cell mean nuclear speckle FISH signal. Boxplots of the same data are shown with an indicator for the median. Asterisks indicate significance based on one-sided Wilcoxon signed-rank tests (****p <= 0.0001, ns: p > 0.05). Scale bar corresponds to 10 µm. **(H)** Most transcripts included in smFISH experiments show a decrease of cytoplasmic RNA intensity and an increase in nuclear to cytoplasmic RNA intensity under DYRK3 inhibition by GSK626616. Each point represents a single gene, points highlighted in red represent genes predicted by APEX2-seq to be depleted from nuclear speckles upon DYRK3 inhibition. **(I)** Example images and quantification across HeLa cells showing changes in translational rates in response to two inhibitors of the nuclear kinase CLK1 and the DYRK3 inhibitor GSK626616 (compared to cells in serum-free media + DMSO), based on incorporation of the puromycin analog O-Propargyl Puromycin (OPP) into nascent peptides.

**Supplementary Figure 5.**
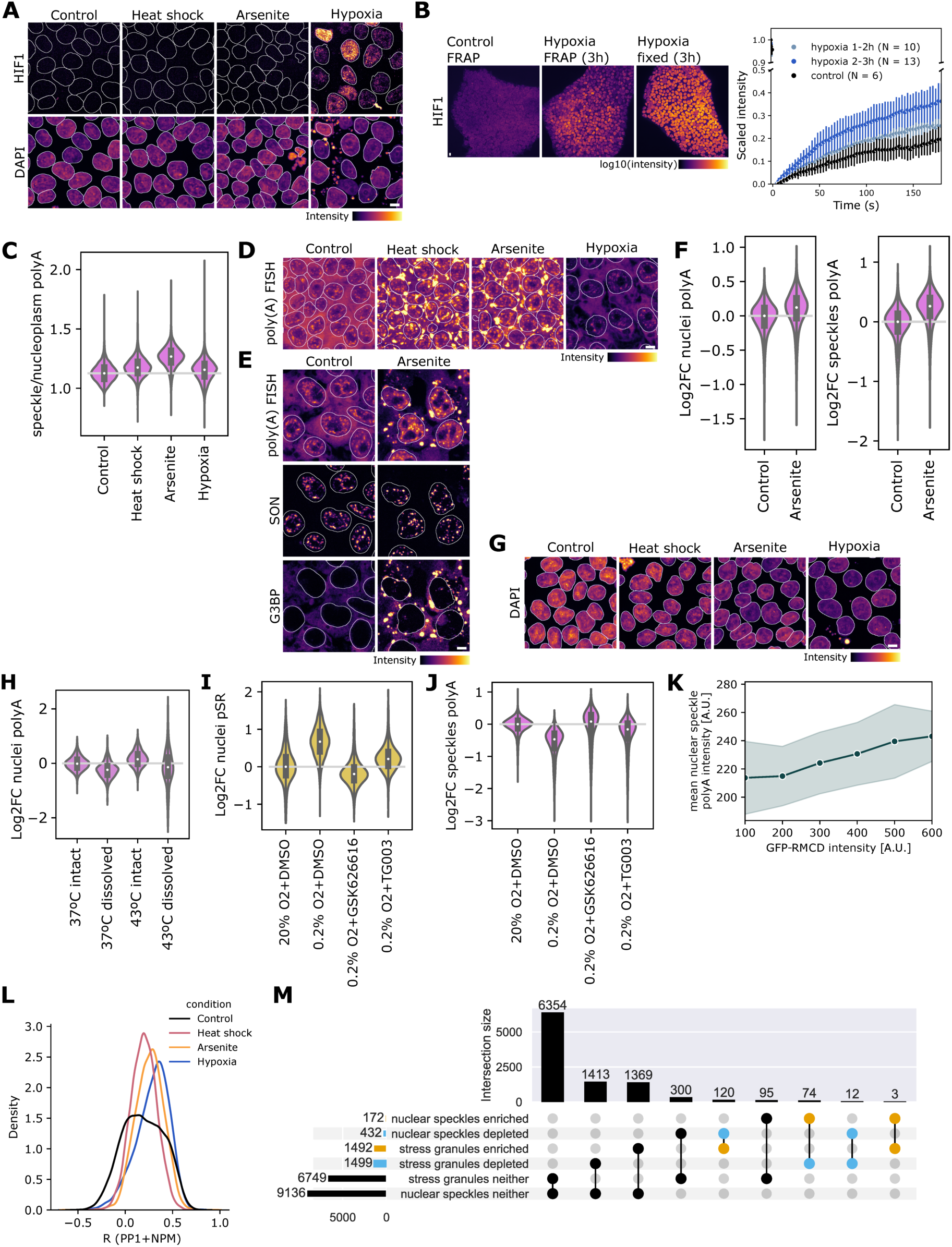
**(A)** Control for induction of cellular response to hypoxia. Example cells show hypoxia-inducible factor-1α (HIF-1α) stabilization and increased expression under hypoxic conditions used (0.2% O2 for 24 hours), but not with heat shock or arsenite-induced oxidative stress. **(B)** HIF-1α staining (left panel) after FRAP (right panel) following brief exposure to hypoxia (0.2% O2 for 1-3 hours for FRAP with fixation at 3 hours) shows a decrease in HIF-1α levels during imaging compared to cells kept and fixed in the hypoxic workstation, but maintenance of higher HIF-1α compared to control cells kept at 20% O2. **(C)** Alternative metric of mRNA retention comparing nuclear speckle to nucleoplasmic intensity of polyA FISH staining shows higher retention within nuclear speckles under arsenite treatment than in other conditions. **(D)** Example images of polyA FISH under environmental perturbations shown in Figure 5E, with mRNA accumulation in stress granules shown. **(E)** Example images of polyA FISH under control and oxidative stress conditions from another experiment where hiPSCs were co-stained for stress granule marker G3BP. **(F)** Quantification of nuclear and nuclear speckle polyA FISH intensity for data shown in (E). **(G)** DAPI staining for cells shown in Fig 5E. **(H)** PolyA FISH intensity in the nucleus in HeLa TREx cells increases with heat shock but decreases with dissolution of nuclear speckles through GFP-CLK1 overexpression. Nuclear speckles were categorized as intact (speckle/nuclear area per cell > 0.1) or dissolved (speckle/nuclear area per cell < 0.05). Log2 fold change compared to the median nuclear polyA RNA in control cells with intact speckles. **(I)** Phospho-SR staining intensity (log2 fold change compared to the DMSO control at 20% O2) under hypoxia (0.2% O2, 24 hours) shows a reduction in pSR with inhibition of DYRK3 by GSK626616 or CLK1 with TG003 to control levels. **(J)** For the data shown in (**I**), polyA FISH signal within nuclear speckles increases under hypoxia with inhibition of DYRK3 or CLK1. **(K)** Expression of a positively charged mixed charge domain during hypoxia recovers mRNA retention within nuclear speckles, depending on the expression level of the mixed charge domain. **(L)** Shifts in pixel correlation between PP1 and a nucleolar marker NPM1 under environmental perturbations. **(M)** An UpSet plot suggests mutual exclusivity between the nuclear speckle and stress granule transcriptomes. Overlaps between enriched (yellow) and depleted (blue) sets of genes in each compartment are highlighted. Scale bars correspond to 10 µM.

## Notes

### Summary of Updates

Figure 4 and Supplementary Figure 4 show additional data, general corrections included.

## References

Bahar Halpern, K., Caspi, I., Lemze, D., Levy, M., Landen, S., Elinav, E., Ulitsky, I., & Itzkovitz, S. (2015). Nuclear Retention of mRNA in Mammalian Tissues. Cell Reports, 13(12), 2653–2662. 10.1016/j.celrep.2015.11.036

Barutcu, A. R., Wu, M., Braunschweig, U., Dyakov, B. J. A., Luo, Z., Turner, K. M., Durbic, T., Lin, Z. Y., Weatheritt, R. J., Maass, P. G., Gingras, A. C., & Blencowe, B. J. (2022). Systematic mapping of nuclear domain-associated transcripts reveals speckles and lamina as hubs of functionally distinct retained introns. Molecular Cell, 82(5), 1035–1052.e9. 10.1016/j.molcel.2021.12.010

Beliveau, B. J., Kishi, J. Y., Nir, G., Sasaki, H. M., Saka, S. K., Nguyen, S. C., Wu, C. Ting, & Yin, P. (2018). OligoMiner provides a rapid, flexible environment for the design of genome-scale oligonucleotide in situ hybridization probes. Proceedings of the National Academy of Sciences of the United States of America, 115(10), E2183–E2192. 10.1073/PNAS.1714530115/SUPPL_FILE/PNAS.1714530115.SD02.PDF

Benoit Bouvrette, L. P., Bovaird, S., Blanchette, M., & Lécuyer, E. (2020). oRNAment: a database of putative RNA binding protein target sites in the transcriptomes of model species. Nucleic Acids Research, 48(D1), D166–D173. 10.1093/NAR/GKZ986

Berg, S., Kutra, D., Kroeger, T., Straehle, C. N., Kausler, B. X., Haubold, C., Schiegg, M., Ales, J., Beier, T., Rudy, M., Eren, K., Cervantes, J. I., Xu, B., Beuttenmueller, F., Wolny, A., Zhang, C., Koethe, U., Hamprecht, F. A., & Kreshuk, A. (2019). ilastik: interactive machine learning for (bio)image analysis. Nature Methods 2019 16:12, 16(12), 1226–1232. 10.1038/s41592-019-0582-9

Berry, S., Müller, M., Rai, A., & Pelkmans, L. (2022). Feedback from nuclear RNA on transcription promotes robust RNA concentration homeostasis in human cells. Cell Systems, 13(6), 454–470.e15. 10.1016/J.CELS.2022.04.005

Bertolotti, A. (2018). The split protein phosphatase system. Biochemical Journal, 475(23), 3707–3723. 10.1042/BCJ20170726

Bolger, A. M., Lohse, M., & Usadel, B. (2014). Trimmomatic: a flexible trimmer for Illumina sequence data. Bioinformatics, 30(15), 2114–2120. 10.1093/BIOINFORMATICS/BTU170

Bollen, M., Peti, W., Ragusa, M. J., & Beullens, M. (2010). The extended PP1 toolkit: Designed to create specificity. Trends in Biochemical Sciences, 35(8), 450–458. 10.1016/j.tibs.2010.03.002

Boutz, P. L., Bhutkar, A., & Sharp, P. A. (2015). Detained introns are a novel, widespread class of post-transcriptionally spliced introns. Genes & Development, 29(1), 63–80. 10.1101/GAD.247361.114

Cao, W., Jamison, S. F., & Garcia-Blanco, M. A. (1997). Both phosphorylation and dephosphorylation of ASF/SF2 are required for pre-mRNA splicing in vitro. RNA, 3(12), 1456–1467. https://rnajournal.cshlp.org/content/3/12/1456.short

Colwill, K., Pawson, T., Andrews, B., Prasad, J., Manley, J. L., Bell, J. C., & Duncan, P. I. (1996). The Clk/Sty protein kinase phosphorylates SR splicing factors and regulates their intranuclear distribution. The EMBO Journal, 15(2), 265. 10.1002/j.1460-2075.1996.tb00357.x

Cox, J., & Mann, M. (2008). MaxQuant enables high peptide identification rates, individualized p.p.b.-range mass accuracies and proteome-wide protein quantification. Nature Biotechnology 2008 26:12, 26(12), 1367–1372. 10.1038/nbt.1511

de Oliveira Freitas Machado, C., Schafranek, M., Brüggemann, M., Cañás, M. C. H., Keller, M., Liddo, A. Di, Brezski, A., Blümel, N., Arnold, B., Bremm, A., Wittig, I., Jaé, N., McNicoll, F., Dimmeler, S., Zarnack, K., & Müller-McNicoll, M. (2023). Poison cassette exon splicing of SRSF6 regulates nuclear speckle dispersal and the response to hypoxia. Nucleic Acids Research, 51(2), 870–890. 10.1093/NAR/GKAC1225

Dias, A. P., Dufu, K., Lei, H., & Reed, R. (2010). A role for TREX components in the release of spliced mRNA from nuclear speckle domains. Nature Communications 2010 1:1, 1(1), 1–10. 10.1038/ncomms1103

Dobin, A., Davis, C. A., Schlesinger, F., Drenkow, J., Zaleski, C., Jha, S., Batut, P., Chaisson, M., & Gingeras, T. R. (2013). STAR: ultrafast universal RNA-seq aligner. Bioinformatics, 29(1), 15. 10.1093/BIOINFORMATICS/BTS635

Fazal, F. M., Han, S., Parker, K. R., Kaewsapsak, P., Xu, J., Boettiger, A. N., Chang, H. Y., & Ting, A. Y. (2019). Atlas of Subcellular RNA Localization Revealed by APEX-Seq. Cell, 178(2), 473–490.e26. 10.1016/j.cell.2019.05.027

Fei, J., Jadaliha, M., Harmon, T. S., Li, I. T. S., Hua, B., Hao, Q., Holehouse, A. S., Reyer, M., Sun, Q., Freier, S. M., Pappu, R. V., Prasanth, K. V., & Ha, T. (2017). Quantitative analysis of multilayer organization of proteins and RNA in nuclear speckles at super resolution. Journal of Cell Science, 130(24), 4180–4192. 10.1242/JCS.206854/VIDEO-1

Galganski, L., Urbanek, M. O., & Krzyzosiak, W. J. (2017). Nuclear speckles: molecular organization, biological function and role in disease. Nucleic Acids Research, 45(18), 10350–10368. 10.1093/NAR/GKX759

Girard, C., Will, C. L., Peng, J., Makarov, E. M., Kastner, B., Lemm, I., Urlaub, H., Hartmuth, K., Luhrmann, R., & Lührmann, R. (2012). Post-transcriptional spliceosomes are retained in nuclear speckles until splicing completion. Nature Communications 2012 3:1, 3(1), 1–12. 10.1038/ncomms1998

Greig, J. A., Nguyen, T. A., Lee, M., Holehouse, A. S., Posey, A. E., Pappu, R. V., & Jedd, G. (2020). Arginine-Enriched Mixed-Charge Domains Provide Cohesion for Nuclear Speckle Condensation. Molecular Cell, 77(6), 1237–1250.e4. 10.1016/j.molcel.2020.01.025

Haltenhof, T., Kotte, A., Bortoli, F. De, Schiefer, S., Meinke, S., Emmerichs, A. K., Petermann, K. K., Timmermann, B., Imhof, P., Franz, A., Loll, B., Wahl, M. C., Preußner, M., & Heyd, F. (2020). A Conserved Kinase-Based Body-Temperature Sensor Globally Controls Alternative Splicing and Gene Expression. Molecular Cell, 78(1), 57–69.e4. 10.1016/J.MOLCEL.2020.01.028

Han, H., Braunschweig, U., Gonatopoulos-Pournatzis, T., Weatheritt, R. J., Hirsch, C. L., Ha, K. C. H., Radovani, E., Nabeel-Shah, S., Sterne-Weiler, T., Wang, J., O’Hanlon, D., Pan, Q., Ray, D., Zheng, H., Vizeacoumar, F., Datti, A., Magomedova, L., Cummins, C. L., Hughes, T. R., … Blencowe, B. J. (2017). Multilayered Control of Alternative Splicing Regulatory Networks by Transcription Factors. Molecular Cell, 65(3), 539–553.e7. 10.1016/J.MOLCEL.2017.01.011

Haupt, A., Grancharova, T., Arakaki, J., Fuqua, M. A., Roberts, B., & Gunawardane, R. N. (2018). Endogenous protein tagging in human induced pluripotent stem cells using CRISPR/Cas9. Journal of Visualized Experiments, 2018(138), e58130. 10.3791/58130

Hershberg, E. A., Camplisson, C. K., Close, J. L., Attar, S., Chern, R., Liu, Y., Akilesh, S., Nicovich, P. R., & Beliveau, B. J. (2021). PaintSHOP enables the interactive design of transcriptome- and genome-scale oligonucleotide FISH experiments. Nature Methods 2021 18:8, 18(8), 937–944. 10.1038/s41592-021-01187-3

Hochberg-Laufer, H., Neufeld, N., Brody, Y., Nadav-Eliyahu, S., Ben-Yishay, R., & Shav-Tal, Y. (2019). Availability of splicing factors in the nucleoplasm can regulate the release of mRNA from the gene after transcription. PLOS Genetics, 15(11), e1008459. 10.1371/JOURNAL.PGEN.1008459

Hoermann, B., Kokot, T., Helm, D., Heinzlmeir, S., Chojnacki, J. E., Schubert, T., Ludwig, C., Berteotti, A., Kurzawa, N., Kuster, B., Savitski, M. M., & Köhn, M. (2020). Dissecting the sequence determinants for dephosphorylation by the catalytic subunits of phosphatases PP1 and PP2A. Nature Communications 2020 11:1, 11(1), 1–20. 10.1038/s41467-020-17334-x

Hornbeck, P. V., Zhang, B., Murray, B., Kornhauser, J. M., Latham, V., & Skrzypek, E. (2015). PhosphoSitePlus, 2014: mutations, PTMs and recalibrations. Nucleic Acids Research, 43(D1), D512–D520. 10.1093/NAR/GKU1267

Hu, Y., Plutz, M., & Belmont, A. S. (2010). Hsp70 gene association with nuclear speckles is Hsp70 promoter specific. Journal of Cell Biology, 191(4), 711–719. 10.1083/JCB.201004041

Hung, C. L. K., Maiuri, T., Bowie, L. E., Gotesman, R., Son, S., Falcone, M., Giordano, J. V., Gillis, T., Mattis, V., Lau, T., Kwan, V., Wheeler, V., Schertzer, J., Singh, K., & Truant, R. (2018). A patient-derived cellular model for Huntington’s disease reveals phenotypes at clinically relevant CAG lengths. Molecular Biology of the Cell, 29(23), 2809–2820. 10.1091/MBC.E18-09-0590/ASSET/IMAGES/LARGE/MBC-29-2809-G004.JPEG

Hutchinson, J. N., Ensminger, A. W., Clemson, C. M., Lynch, C. R., Lawrence, J. B., & Chess, A. (2007). A screen for nuclear transcripts identifies two linked noncoding RNAs associated with SC35 splicing domains. BMC Genomics, 8(1), 1–16. 10.1186/1471-2164-8-39/FIGURES/6

Ilık, İ. A., Malszycki, M., Lübke, A. K., Schade, C., Meierhofer, D., & Aktaş, T. (2020). Son and srrm2 are essential for nuclear speckle formation. ELife, 9, 1–48. 10.7554/ELIFE.60579

Jain, A., & Vale, R. D. (2017). RNA phase transitions in repeat expansion disorders. Nature 2017 546:7657, 546(7657), 243–247. 10.1038/nature22386

Jakubauskiene, E., Vilys, L., Makino, Y., Poellinger, L., & Kanopka, A. (2015). Increased serine-arginine (SR) protein phosphorylation changes pre-mRNA splicing in hypoxia. Journal of Biological Chemistry, 290(29), 18079–18089. 10.1074/jbc.M115.639690

Kaewsapsak, P., Shechner, D. M., Mallard, W., Rinn, J. L., & Ting, A. Y. (2017). Live-cell mapping of organelle-associated RNAs via proximity biotinylation combined with protein-RNA crosslinking. ELife, 6. 10.7554/ELIFE.29224

Kedersha, N., & Anderson, P. (2002). Stress granules: sites of mRNA triage that regulate mRNA stability and translatability. Biochemical Society Transactions, 30(6), 963–969. 10.1042/BST0300963

Khong, A., Matheny, T., Jain, S., Mitchell, S. F., Wheeler, J. R., & Parker, R. (2017). The Stress Granule Transcriptome Reveals Principles of mRNA Accumulation in Stress Granules. Molecular Cell, 68(4), 808–820.e5. 10.1016/J.MOLCEL.2017.10.015

Kim, J., Han, K. Y., Khanna, N., Ha, T., & Belmont, A. S. (2019). Nuclear speckle fusion via long-range directional motion regulates speckle morphology after transcriptional inhibition. Journal of Cell Science, 132(8). 10.1242/JCS.226563/VIDEO-9

Kim, J., Venkata, N. C., Hernandez Gonzalez, G. A., Khanna, N., & Belmont, A. S. (2020). Gene expression amplification by nuclear speckle association. The Journal of Cell Biology, 219(1). 10.1083/jcb.201904046

Kishi, J. Y., Lapan, S. W., Beliveau, B. J., West, E. R., Zhu, A., Sasaki, H. M., Saka, S. K., Wang, Y., Cepko, C. L., & Yin, P. (2019). SABER amplifies FISH: enhanced multiplexed imaging of RNA and DNA in cells and tissues. Nature Methods 2019 16:6, 16(6), 533–544. 10.1038/s41592-019-0404-0

Kucera, M., Isserlin, R., Arkhangorodsky, A., & Bader, G. D. (2016). AutoAnnotate: A Cytoscape app for summarizing networks with semantic annotations. F1000Research 2016 5:1717, 5, 1717. 10.12688/f1000research.9090.1

Lester, E., Ooi, F. K., Bakkar, N., Ayers, J., Woerman, A. L., Wheeler, J., Bowser, R., Carlson, G. A., Prusiner, S. B., & Parker, R. (2021). Tau aggregates are RNA-protein assemblies that mislocalize multiple nuclear speckle components. Neuron, 109(10), 1675–1691.e9. 10.1016/J.NEURON.2021.03.026

Li, H. D., Funk, C. C., & Price, N. D. (2020). IREAD: A tool for intron retention detection from RNA-seq data. BMC Genomics, 21(1), 1–11. 10.1186/S12864-020-6541-0/FIGURES/9

Liao, Y., Smyth, G. K., & Shi, W. (2014). featureCounts: an efficient general purpose program for assigning sequence reads to genomic features. Bioinformatics (Oxford, England), 30(7), 923–930. 10.1093/BIOINFORMATICS/BTT656

Liao, Y., Smyth, G. K., & Shi, W. (2019). The R package Rsubread is easier, faster, cheaper and better for alignment and quantification of RNA sequencing reads. Nucleic Acids Research, 47(8), e47–e47. 10.1093/NAR/GKZ114

Liu, J., Xu, Y., Stoleru, D., & Salic, A. (2012). Imaging protein synthesis in cells and tissues with an alkyne analog of puromycin. Proceedings of the National Academy of Sciences of the United States of America, 109(2), 413–418. 10.1073/PNAS.1111561108/-/DCSUPPLEMENTAL/PNAS.1111561108_SI.PDF

Love, M. I., Huber, W., & Anders, S. (2014). Moderated estimation of fold change and dispersion for RNA-seq data with DESeq2. Genome Biology, 15(12), 1–21. 10.1186/S13059-014-0550-8/FIGURES/9

Lun, X. K., Szklarczyk, D., Gábor, A., Dobberstein, N., Zanotelli, V. R. T., Saez-Rodriguez, J., von Mering, C., & Bodenmiller, B. (2019). Analysis of the Human Kinome and Phosphatome by Mass Cytometry Reveals Overexpression-Induced Effects on Cancer-Related Signaling. Molecular Cell, 74(5), 1086–1102.e5. 10.1016/J.MOLCEL.2019.04.021

Luo, Y., Liu, C., Cerbini, T., San, H., Lin, Y., Chen, G., Rao, M. S., & Zou, J. (2014). Stable enhanced green fluorescent protein expression after differentiation and transplantation of reporter human induced pluripotent stem cells generated by AAVS1 transcription activator-like effector nucleases. Stem Cells Translational Medicine, 3(7), 821–835. 10.5966/SCTM.2013-0212

Maćkiewicz, A., & Ratajczak, W. (1993). Principal components analysis (PCA). Computers & Geosciences, 19(3), 303–342. 10.1016/0098-3004(93)90090-R

Martins, S. B., Rino, J., Carvalho, T., Carvalho, C., Yoshida, M., Klose, J. M., Almeida, S. F. De, & Carmo-Fonseca, M. (2011). Spliceosome assembly is coupled to RNA polymerase II dynamics at the 3ʹ end of human genes. Nature Structural & Molecular Biology 2011 18:10, 18(10), 1115–1123. 10.1038/nsmb.2124

Mateju, D., & Chao, J. A. (2022). Stress granules: regulators or by-products? The FEBS Journal, 289(2), 363–373. 10.1111/FEBS.15821

McInnes, L., Healy, J., & Melville, J. (2018). *UMAP: Uniform Manifold Approximation and Projection for Dimension Reduction*. https://arxiv.org/abs/1802.03426v3

McInnes, L., Healy, J., Saul, N., & Großberger, L. (2018). UMAP: Uniform Manifold Approximation and Projection. Journal of Open Source Software, 3(29), 861. 10.21105/JOSS.00861

Merico, D., Isserlin, R., Stueker, O., Emili, A., & Bader, G. D. (2010). Enrichment map: a network-based method for gene-set enrichment visualization and interpretation. PloS One, 5(11). 10.1371/JOURNAL.PONE.0013984

Mermoud, J. E., Cohen, P. T. W., & Lamond, A. I. (1994). Regulation of mammalian spliceosome assembly by a protein phosphorylation mechanism. The EMBO Journal, 13(23), 5679–5688. 10.1002/J.1460-2075.1994.TB06906.X

Misteli, T., & Spector, D. L. (1996). Serine/threonine phosphatase 1 modulates the subnuclear distribution of pre-mRNA splicing factors. Https://Doi.Org/10.1091/Mbc.7.10.1559, 7(10), 1559–1572. https://doi.org/10.1091/MBC.7.10.1559

Miyagawa, R., Tano, K., Mizuno, R., Nakamura, Y., Ijiri, K., Rakwal, R., Shibato, J., Masuo, Y., Mayeda, A., Hirose, T., & Akimitsu, N. (2012). Identification of cis- and trans-acting factors involved in the localization of MALAT-1 noncoding RNA to nuclear speckles. RNA, 18(4), 738. 10.1261/RNA.028639.111

Moon, S. L., Morisaki, T., Khong, A., Lyon, K., Parker, R., & Stasevich, T. J. (2019). Multicolour single-molecule tracking of mRNA interactions with RNP granules. Nature Cell Biology 2019 21:2, 21(2), 162–168. 10.1038/s41556-018-0263-4

Mootha, V. K., Lindgren, C. M., Eriksson, K. F., Subramanian, A., Sihag, S., Lehar, J., Puigserver, P., Carlsson, E., Ridderstråle, M., Laurila, E., Houstis, N., Daly, M. J., Patterson, N., Mesirov, J. P., Golub, T. R., Tamayo, P., Spiegelman, B., Lander, E. S., Hirschhorn, J. N., … Groop, L. C. (2003). PGC-1α-responsive genes involved in oxidative phosphorylation are coordinately downregulated in human diabetes. Nature Genetics 2003 34:3, 34(3), 267–273. 10.1038/ng1180

Ninomiya, K., Kataoka, N., & Hagiwara, M. (2011). Stress-responsive maturation of Clk1/4 pre-mRNAs promotes phosphorylation of SR splicing factor. Journal of Cell Biology, 195(1), 27–40. 10.1083/JCB.201107093

Oceguera-Yanez, F., Kim, S. Il, Matsumoto, T., Tan, G. W., Xiang, L., Hatani, T., Kondo, T., Ikeya, M., Yoshida, Y., Inoue, H., & Woltjen, K. (2016). Engineering the AAVS1 locus for consistent and scalable transgene expression in human iPSCs and their differentiated derivatives. Methods (San Diego, Calif.), 101, 43–55. 10.1016/J.YMETH.2015.12.012

O’Keefe, R. T., Mayeda, A., Sadowski, C. L., Krainer, A. R., & Spector, D. L. (1994). Disruption of pre-mRNA splicing in vivo results in reorganization of splicing factors. Journal of Cell Biology, 124(3), 249–260. 10.1083/JCB.124.3.249

Padrón, A., Iwasaki, S., & Ingolia, N. T. (2019). Proximity RNA Labeling by APEX-Seq Reveals the Organization of Translation Initiation Complexes and Repressive RNA Granules. Molecular Cell, 75(4), 875–887.e5. 10.1016/j.molcel.2019.07.030

Passaro, M., Martinovic, M., Bevilacqua, V., Hershberg, E. A., Rossetti, G., Beliveau, B. J., Bonnal, R. J. P., & Pagani, M. (2020). OligoMinerApp: a web-server application for the design of genome-scale oligonucleotide in situ hybridization probes through the flexible OligoMiner environment. Nucleic Acids Research, 48(W1), W332–W339. 10.1093/NAR/GKAA251

Patro, R., Duggal, G., Love, M. I., Irizarry, R. A., & Kingsford, C. (2017). Salmon provides fast and bias-aware quantification of transcript expression. Nature Methods 2017 14:4, 14(4), 417–419. 10.1038/nmeth.4197

Paul, S., Arias, M. A., Wen, L., Liao, S. E., Zhang, J., Wang, X., Regev, O., & Fei, J. (2022). Nuclear speckle-localized RNAs exhibit preferential positioning and orientation. BioRxiv, 2022.10.17.512423. 10.1101/2022.10.17.512423

Périer, R. C., Junier, T., & Bucher, P. (1998). The Eukaryotic Promoter Database EPD. Nucleic Acids Research, 26(1), 353–357. 10.1093/NAR/26.1.353

Politz, J. C. R., Tuft, R. A., Prasanth, K. V., Baudendistel, N., Fogarty, K. E., Lifshitz, L. M., Langowski, J., Spector, D. L., & Pederson, T. (2006). Rapid, diffusional shuttling of poly(A) RNA between nuclear speckles and the nucleoplasm. Molecular Biology of the Cell, 17(3), 1239–1249. 10.1091/mbc.E05-10-0952

Rai, A. K., Chen, J. X., Selbach, M., & Pelkmans, L. (2018). Kinase-controlled phase transition of membraneless organelles in mitosis. Nature, 559(7713), 211–216. 10.1038/s41586-018-0279-8

Randolph, L. N., Bao, X., Zhou, C., & Lian, X. (2017). An all-in-one, Tet-On 3G inducible PiggyBac system for human pluripotent stem cells and derivatives. Scientific Reports 2017 7:1, 7(1), 1–8. 10.1038/s41598-017-01684-6

Rino, J., Carvalho, T., Braga, J., Desterro, J. M. P. P., Lührmann, R., & Carmo-Fonseca, M. (2007). A Stochastic View of Spliceosome Assembly and Recycling in the Nucleus. PLOS Computational Biology, 3(10), e201. 10.1371/JOURNAL.PCBI.0030201

Roberts, B., Haupt, A., Tucker, A., Grancharova, T., Arakaki, J., Fuqua, M. A., Nelson, A., Hookway, C., Ludmann, S. A., Mueller, I. A., Yang, R., Horwitz, R., Rafelski, S. M., & Gunawardane, R. N. (2017). Systematic gene tagging using CRISPR/Cas9 in human stem cells to illuminate cell organization. Molecular Biology of the Cell, 28(21), 2854–2874. 10.1091/mbc.E17-03-0209

Saavedra, C., Tuug, K. S., Amberg, D. C., Hopper, A. K., & Cole, C. N. (1996). Regulation of mRNA export in response to stress in Saccharomyces cerevisiae. Genes & Development, 10(13), 1608–1620. 10.1101/GAD.10.13.1608

Shalgi, R., Hurt, J. A., Lindquist, S., & Burge, C. B. (2014). Widespread inhibition of posttranscriptional splicing shapes the cellular transcriptome following heat shock. Cell Reports, 7(5), 1362–1370. 10.1016/j.celrep.2014.04.044

Shannon, P., Markiel, A., Ozier, O., Baliga, N. S., Wang, J. T., Ramage, D., Amin, N., Schwikowski, B., & Ideker, T. (2003). Cytoscape: a software environment for integrated models of biomolecular interaction networks. Genome Research, 13(11), 2498–2504. 10.1101/GR.1239303

Shi, Y., Reddy, B., & Manley, J. L. (2006). PP1/PP2A Phosphatases Are Required for the Second Step of Pre-mRNA Splicing and Target Specific snRNP Proteins. Molecular Cell, 23(6), 819–829. 10.1016/j.molcel.2006.07.022

Shin, C., Feng, Y., & Manley, J. L. (2004). Dephosphorylated SRp38 acts as a splicing repressor in response to heat shock. Nature 2004 427:6974, 427(6974), 553–558. 10.1038/nature02288

Smalec, B. M., Ietswaart, R., Choquet, K., McShane, E., West, E. R., & Churchman, L. S. (2022). Genome-wide quantification of RNA flow across subcellular compartments reveals determinants of the mammalian transcript life cycle. BioRxiv, 2022.08.21.504696. 10.1101/2022.08.21.504696

Smith, K. P., Moen, P. T., Wydner, K. L., Coleman, J. R., & Lawrence, J. B. (1999). Processing of endogenous pre-mRNAs in association with SC-35 domains is gene specific. Journal of Cell Biology, 144(4), 617–629. 10.1083/jcb.144.4.617

Söding, J., Zwicker, D., Sohrabi-Jahromi, S., Boehning, M., & Kirschbaum, J. (2020). Mechanisms for Active Regulation of Biomolecular Condensates. Trends in Cell Biology, 30(1), 4–14. 10.1016/j.tcb.2019.10.006

Sommer, C., Straehle, C., Kothe, U., & Hamprecht, F. A. (2011). Ilastik: Interactive learning and segmentation toolkit. Proceedings - International Symposium on Biomedical Imaging, 230–233. 10.1109/ISBI.2011.5872394

Stringer, C., Wang, T., Michaelos, M., & Pachitariu, M. (2020). Cellpose: a generalist algorithm for cellular segmentation. Nature Methods 2020 18:1, 18(1), 100–106. 10.1038/s41592-020-01018-x

Subramanian, A., Tamayo, P., Mootha, V. K., Mukherjee, S., Ebert, B. L., Gillette, M. A., Paulovich, A., Pomeroy, S. L., Golub, T. R., Lander, E. S., & Mesirov, J. P. (2005). Gene set enrichment analysis: A knowledge-based approach for interpreting genome-wide expression profiles. Proceedings of the National Academy of Sciences of the United States of America, 102(43), 15545–15550. 10.1073/PNAS.0506580102/SUPPL_FILE/06580FIG7.JPG

Tapial, J., Ha, K. C. H., Sterne-Weiler, T., Gohr, A., Braunschweig, U., Hermoso-Pulido, A., Quesnel-Vallières, M., Permanyer, J., Sodaei, R., Marquez, Y., Cozzuto, L., Wang, X., Gómez-Velázquez, M., Rayon, T., Manzanares, M., Ponomarenko, J., Blencowe, B. J., & Irimia, M. (2017). An atlas of alternative splicing profiles and functional associations reveals new regulatory programs and genes that simultaneously express multiple major isoforms. Genome Research, 27(10), 1759–1768. 10.1101/GR.220962.117

Trinkle-Mulcahy, L., Ajuh, P., Prescott, A., Claverie-Martin, F., Cohen, S., Lamond, A. I., & Cohen, P. (1999). Nuclear organisation of NIPP1, a regulatory subunit of protein phosphatase 1 that associates with pre-mRNA splicing factors. Journal of Cell Science, 112(2), 157–168. 10.1242/JCS.112.2.157

Trinkle-Mulcahy, L., Sleeman, J. E., & Lamond, A. I. (2001). Dynamic targeting of protein phosphatase 1 within the nuclei of living mammalian cells. Journal of Cell Science, 114(23), 4219–4228. 10.1242/JCS.114.23.4219

Türker, C., Akal, F., Joho, D., Panse, C., Barkow-Oesterreicher, S., Rehrauer, H., & Schlapbach, R. (2010). B-fabric: The Swiss army knife for life sciences. Advances in Database Technology - EDBT 2010 - 13th International Conference on Extending Database Technology, Proceedings, 717–720. 10.1145/1739041.1739135

Twyffels, L., Gueydan, C., & Kruys, V. (2011). Shuttling SR proteins: more than splicing factors. The FEBS Journal, 278(18), 3246–3255. 10.1111/J.1742-4658.2011.08274.X

Vaquero-Garcia, J., Barrera, A., Gazzara, M. R., Gonzalez-Vallinas, J., Lahens, N. F., Hogenesch, J. B., Lynch, K. W., & Barash, Y. (2016). A new view of transcriptome complexity and regulation through the lens of local splicing variations. ELife, 5(FEBRUARY2016). 10.7554/ELIFE.11752

Wang, J. T., Smith, J., Chen, B. C., Schmidt, H., Rasoloson, D., Paix, A., Lambrus, B. G., Calidas, D., Betzig, E., & Seydoux, G. (2014). Regulation of RNA granule dynamics by phosphorylation of serine-rich, intrinsically disordered proteins in C. elegans. ELife, 3. 10.7554/ELIFE.04591

Wang, K., Wang, L., Wang, J., Chen, S., Shi, M., & Cheng, H. (2018). Intronless mRNAs transit through nuclear speckles to gain export competence. Journal of Cell Biology, 217(11), 3912–3929. 10.1083/jcb.201801184

Wegener, M., & Müller-McNicoll, M. (2018). Nuclear retention of mRNAs – quality control, gene regulation and human disease. Seminars in Cell & Developmental Biology, 79, 131–142. 10.1016/J.SEMCDB.2017.11.001

Wippich, F., Bodenmiller, B., Trajkovska, M. G., Wanka, S., Aebersold, R., & Pelkmans, L. (2013). Dual Specificity Kinase DYRK3 Couples Stress Granule Condensation/Dissolution to mTORC1 Signaling. Cell, 152(4), 791–805. 10.1016/J.CELL.2013.01.033

Wu, D., Wever, V. De, Derua, R., Winkler, C., Beullens, M., Eynde, A. Van, & Bollen, M. (2018). A substrate-trapping strategy for protein phosphatase PP1 holoenzymes using hypoactive subunit fusions. Journal of Biological Chemistry, 293(39), 15152–15162. 10.1074/jbc.RA118.004132

Xu, Q., Schlabach, M. R., Hannon, G. J., & Elledge, S. J. (2009). Design of 240,000 orthogonal 25mer DNA barcode probes. Proceedings of the National Academy of Sciences of the United States of America, 106(7), 2289–2294. 10.1073/PNAS.0812506106/ASSET/64B745AE-1AE3-4991-957F-3109B3830384/ASSETS/GRAPHIC/ZPQ9990965120003.JPEG

Yusa, K., Zhou, L., Li, M. A., Bradley, A., & Craig, N. L. (2011). A hyperactive piggyBac transposase for mammalian applications. Proceedings of the National Academy of Sciences of the United States of America, 108(4), 1531–1536. 10.1073/PNAS.1008322108/SUPPL_FILE/PNAS.201008322SI.PDF

Zander, G., Hackmann, A., Bender, L., Becker, D., Lingner, T., Salinas, G., & Krebber, H. (2016). mRNA quality control is bypassed for immediate export of stress-responsive transcripts. Nature 2016 540:7634, 540(7634), 593–596. 10.1038/nature20572

Zhang, B., Gunawardane, L., Niazi, F., Jahanbani, F., Chen, X., & Valadkhan, S. (2014). A Novel RNA Motif Mediates the Strict Nuclear Localization of a Long Noncoding RNA. Molecular and Cellular Biology, 34(12), 2318–2329. 10.1128/MCB.01673-13/SUPPL_FILE/TMCB_A_12275259_SM0001.PDF

Zhang, L., Zhang, Y., Chen, Y., Gholamalamdari, O., Wang, Y., Ma, J., & Belmont, A. S. (2021). TSA-seq reveals a largely conserved genome organization relative to nuclear speckles with small position changes tightly correlated with gene expression changes. Genome Research, 31(2), 251–264. 10.1101/GR.266239.120/-/DC1

Zhang, Q., Kota, K. P., Alam, S. G., Nickerson, J. A., Dickinson, R. B., & Lele, T. P. (2016). Coordinated Dynamics of RNA Splicing Speckles in the Nucleus. Journal of Cellular Physiology, 231(6), 1269–1275. 10.1002/JCP.25224

Zhu, A., Srivastava, A., Ibrahim, J. G., Patro, R., & Love, M. I. (2019). Nonparametric expression analysis using inferential replicate counts. Nucleic Acids Research, 47(18), e105–e105. 10.1093/NAR/GKZ622

