## Supplementary material for "Phosphorylation-controlled cohesion of a nuclear condensate regulates mRNA retention": Readme

supplementary_table_1.csv:
Long format table that contains DESeq2 results of different comparisons. The descriptor column indicates which comparison the row data comes from. The descriptor can have the following values:

fractionation_chr_vs_cyt: Result of fractionation sequencing experiment comparing chromatin to cytoplasm fraction
fractionation_chr_vs_nuc: Result of fractionation sequencing experiment comparing chromatin to nuclei fraction
fractionation_nuc_vs_cyt: Result of fractionation sequencing experiment comparing nuclei to cytoplasm fraction
speckle_enrichment_input: Result of APEX2seq experiment comparing input samples of APEX2-SON to APEX2-NLS cell line
speckle_enrichment_wo_input_normalization: Result of APEX2-seq experiment comparing biontinylated samples of APEX2-SON to APEX2-NLS cell line (without input normalization)
speckle_enrichment_lm_residual: Result of APEX2-seq experiment comparing biotinylated samples of APEX2-SON to APEX2-NLS cell line & normalizing for changes in input samples with a linear model
speckle_enrichment_GSK_DMSO: Result of APEX2-seq experiment comparing change in speckle enrichment in GSK626616 treated samples vs. DMSO treated samples (using interaction term treatment:cell_line)
speckle_enrichment_DRB_DMSO: Result of APEX2-seq experiment comparing change in speckle enrichment in DRB treated samples vs. DMSO treated samples (using interaction term treatment:cell_line)
speckle_enrichment_PLAD_DMSO: Result of APEX2-seq experiment comparing change in speckle enrichment in Pladienolide-B treated samples vs. DMSO treated samples (using interaction term treatment:cell_line)


supplementary_table_2.csv:
Metadata of smFISH probes designed to validate APEX2-seq results. Columns are:

gene_name: names of target genes
color: color of the probes
sequence: raw sequences as generated from paintshop or oligominer
pool_name: name of the pool in which they were ordered
sequence_with_barcodes: probe sequence after extending with color-specific barcodes


supplementary_table_3.csv:
Table containing metadata for all transcript features used for training of the multilinear regression model and UMAP embedding. Columns are:

feature name: name of the feature
correlation with transcript log2FC nuclear speckle enrichment: Pearson correlation coefficient with transcript log2FC nuclear speckle enrichment after preprocessing of data
frequency of occurence in top 10 loadings of top 10 PCs: frequency at which the feature appears in all features of the top 10 loadings of top 10 pcs across 100 iterations of model training (a frequency of 1 means it appears in the top10 loadings of each of the top 10 PCs).
feature category: category of the feature; can be general sequence feature, RBP binding motif, rate feature, promotor motif or transcript expression level (TPM)
source: source from which the feature values originate

supplementary_table_4.csv:
Table containing information about smFISH branching probes used for signal amplification. Columns are:

probe_name: name of the probe, can be secondary, tertiary, quaternary or label probe. Probes were used in smFISH in this order to amplify signal from gene-specific primary probes.
color: color of the fluorophor the probe ultimately encodes (which label probe finally is bound)
sequence: sequence of the probe
length: length of the probe
branching_motif: motif that is present in the probe and allows the next level of probes to bind
number_of_branching_motifs: number of branching motifs present in the sequence of the probe
binds_to: sequence to which the probe will bind (a sequence present in the branching probe of the previous level)
